# A structure-guided pipeline yields peptide inhibitors that disarm fungal peptidase-driven virulence and resistance

**DOI:** 10.64898/2026.04.10.717560

**Authors:** Davier Gutierrez-Gongora, Jacob Hambly, Norris Chan, Oscar Romero, Michael Woods, Esther Olabisi-Adeniyi, Aimee Dawe, Maria Juliana Mantilla, Gregory A. Wasney, Jared Deyarmin, Stephanie N. Samra, Pedro A. Valiente, Melanie Alpaugh, Adnane Sellam, Ryan S. Prosser, Andrew Hamilton-Wright, Jennifer Geddes-McAlister

## Abstract

Fungal infections are a major global health challenge, with current antifungal therapies limited by toxicity, cost, and resistance. For *Cryptococcus neoformans*, key virulence factors that initiate and sustain infection are regulated by fungal peptidases to produce a polysaccharide capsule, promote immune evasion, and support antifungal resistance. These peptidases represent promising targets for antivirulent therapeutic strategies. Here, we developed a computational pipeline to predict and design peptide- and protein-based inhibitors against cryptococcal peptidases. Specifically, we targeted three virulence-associated peptidases: Rim13 (cysteine), May1 (aspartic), and CnMpr1 (metallo). Cysteine peptidase inhibition decreased capsule/cell size ratios without impeding fungal growth and reduced fungal survival within macrophages. Similarly, aspartic peptidase inhibition enhanced fungal clearance within alveolar macrophages and disrupted biofilm formation with additive effects towards fluconazole susceptibility in resistant strains. Additionally, metallopeptidase inhibition through catalytic zinc chelation and blocked substrate binding led to enhanced enzymatic inhibition and reduced *in vitro* blood-brain barrier crossing. Moreover, an *in vivo* larval model assessing inhibitor efficacy produced additive effects with fluconazole and lacked host cell cytotoxicity and fungicidal properties, reinforcing anti-virulence mechanisms and therapeutic potential while limiting the evolution of resistance. Further, global proteome profiling of inhibitor treated cells defined a mechanism of cell wall disruption, impeding fungal virulence. Taken together, the designed peptidase inhibitors exhibited potent antifungal activity without harming mammalian cells, establishing a predictive framework for rational scaffold design of next-generation antifungals that disarm the pathogen enabling immune-mediated clearance.

## Introduction

Fungal pathogens present a critical health threat with rising incidence and mortality rates. For the globally relevant opportunistic human fungal pathogen, *Cryptococcus neoformans,* infections primarily affect immunocompromised individuals, causing approximately 20% of HIV/AIDS related deaths worldwide ^1,2^. Gold standard treatment options for cryptococcal infections require hospitalization due to severe health implications or prolonged (>6 months) exposure that promote the evolution of resistance and reduced efficacy ^3,4^. Additionally, similarity between biochemical pathways and structures of fungal and mammalian cells confounds the challenge of developing new, safe antifungal agents ^5^, demanding alternative approaches that disarm the pathogen without harming the host.

The ability of *C. neoformans* to cause disease and the severity of infection depends on the production of crucial virulence factors, including i) thermotolerance (i.e., capacity to grow at host body temperature) ^6^, ii) polysaccharide capsule (i.e., immune system evasion and modulation) ^7^, iii) melanin (i.e., protection against stress and antifungal drugs) ^7,8^ and, iv) extracellular enzymes (i.e., host tissue invasion and damage) ^9^. Regulation of these virulence factors is controlled by peptidases, belonging to the metallo, cysteine, and aspartic families^10^. For instance, Rim13 (CNAG_05601), a C2 calpain-like cysteine peptidase, interacts with the RIM pathway to promote adaptation to alkaline pH, thereby strengthening the fungal cell wall and capsule attachment to evade the immune system^11^. Similarly, May1 (CNAG_05872), an A1 pepsin-like aspartic peptidase, is secreted for survival in acidic environments, such as the macrophage phagolysosome ^10,11^, and CnMpr1 (CNAG_04735), a M36 metallopeptidase, is secreted by *C. neoformans* near the blood-brain barrier (BBB) to promote crossing and invasion^12^. Using virulence factors as a therapeutic target offers several advantages, including the possibility of disarming the pathogen, thereby reducing the rate of antifungal resistance by imposing lower selective pressure than killing the pathogen directly, and to promote more efficient clearance by the immune system ^9,13,14^. Importantly, inhibition of cryptococcal peptidase is a promising treatment option for fungal infections ^15,16^, and highlights the potential of peptidase inhibitors as antifungal agents ^17,18^.

In nature, peptidase inhibitors vary in size and chemical structure (e.g., proteins, peptides or metabolites) with specificity towards one or more families of peptidases and diverse mechanisms of inhibition ^19,20^. For competitive peptidase inhibitors, which mimic the natural substrates of peptidases but resist (e.g., irreversible) or slow (e.g., reversible) cleavage, thereby, blocking the peptidase catalytic cycle, deliver high specificity, reduced off-target effects and increased therapeutic efficacy ^21^. Thus, due to their high affinity for the target peptidase, naturally-derived inhibitors can be highly potent, requiring lower doses to achieve the desired effect. We previously reported the discovery of peptidase inhibitors from natural sources (i.e., mollusks) with antifungal effects towards *C. neoformans*, demonstrating the promise of naturally sourced inhibitors against critical fungal virulence regulators ^22,23^. However, these untargeted studies lacked fungal peptidase specificity and *a priori* knowledge of prospective inhibitors derived from the mollusk extracts, limiting the discovery power of peptidase-specific inhibitors. Despite these limitations, promise for the discovery of peptidase inhibitors with specific and potent antifungal activity is possible with computational engineering using structure-based rational design.

In this study, we curated a peptidase-inhibitor complex database and developed an AlphaFold-Multimer (AFM)-based pipeline^24–26^, integrating Alpha-Pulldown^26^ with AlphaFold2^27^ for multichain protein complex prediction to model protein-protein interactions, including enzyme-inhibitor interactions^28^. Using this AFM-based pipeline, we designed and prioritized eight peptide-based inhibitors that target virulence-related peptidases of *C. neoformans*: i) Rim13; involved in immune system evasion through capsule production^29,30^, ii) May1; involved in macrophage clearance survival in acidic compartments^31^ and, iii) CnMpr1; critical for fungal cell dissemination and BBB crossing^12^. *In vitro* assessment of the computationally designed peptidase inhibitors revealed function-specific disruption of fungal virulence, complemented by enhanced macrophage fungal clearance, re-sensitization to fluconazole, impaired crossing of the BBB, and additive effects with fluconazole to reduce virulence *in vivo* using an invertebrate model of cryptococcal infection. These data were complemented with proteome profiling to assess global alterations upon inhibitor treatment and propose a mechanism of action by cell wall disruption for reduced virulence. Collectively, we establish a computational framework integrated with *in vitro* and *in vivo* characterization to design and validate fungal peptidase inhibitors, advancing a new class of immune-supportive antifungal therapeutics that disable pathogenic mechanisms without compromising host integrity.

## Materials and Methods

### Peptidase-inhibitor screening pipeline

A custom end-to-end pipeline was created (https://github.com/HamSlice7/Peptidase-Inhibitor-Screening-Pipeline) using AFM. Three inputs were required for each run: 1) a FASTA file containing the amino acid sequence of the target peptidase, 2) a FASTA file containing amino acid sequences of inhibitor candidates, and 3) the residue number of the peptidase active site. Outputs included scoring metrics: interface predicted template modeling (ipTM), localized interaction score (LIS), and localized interaction area (LIA) ^25,28^. These scores were used to predict inhibitors that interact with a peptidase drug target. Additionally, the ratio of the solvent accessibility surface area at the active site (rSASA) and the distance of the inhibitor to the active site were used to predict competitive inhibitors ^32^. If the active site residue of the peptidase was unknown, the parameter was set to ‘0’ (i.e., rSASA and distance to the active site scores were removed) for the predicted peptidase-inhibitor complexes. All scoring results for each predicted complex were compiled for downstream analysis and interpretation.

### Preparing the testing dataset

Experimentally validated 108 peptidase-inhibitor pairs were gathered from the Protein Databank (PDB; https://www.rcsb.org/). Each peptidase belonged to one of four main peptidase families: aspartic, serine, metallo, or cysteine. The dataset was split into two: peptide or protein inhibitors, based on amino acid sequence length. Performance of AFM to classify true positive (TP) and true negative (TN) peptidase-inhibitor pairs included 62 peptidase-inhibitor pairs with inhibitors < 30 amino acids (i.e., peptide inhibitors) and 46 peptidase-inhibitor pairs with inhibitors > 30 amino acids (i.e., protein inhibitors). Next, for each data set, each peptidase was randomly matched with an inhibitor from a different family, creating a set of TN peptidase-inhibitor pairs not expected to interact. This resulted in 124 and 92 peptidase-inhibitor pairs for the peptide and protein inhibitor datasets, respectfully. Next, amino acid sequences of the peptidases and the inhibitors from these datasets were built into two separate FASTA files. The signal peptide and propeptide sequences (as designated within the PDB) were removed to provide the mature peptidase form.

### Running the testing dataset

The FASTA files were transferred to a high-performance computing cluster (HPCC) and the custom peptidase-inhibitor pipeline was cloned from GitHub (https://github.com/HamSlice7/Peptidase-Inhibitor-Screening-Pipeline/tree/main/pipeline_custom). The custom pipeline operated Alpha-Pulldown in ‘custom mode’ with a text file of peptidase-inhibitor pairs from the protein and peptide inhibitor data set used as an additional input. No active site sequence of the peptidases were provided.

### Assessing the testing dataset

The run output included: i) peptidase-peptide inhibitor testing dataset, and ii) peptidase-protein inhibitor testing dataset. By default, five structure predictions of each peptidase-inhibitor pair were generated for AFM conformational testing. Binary labels were added to each peptidase-inhibitor pair based on known (PDB) or random matching. The mean and max ipTM, LIS and LIA from the five predicted structures for each peptidase-inhibitor pair were extracted. A Mann-Whitney U test was performed for each scoring metric in each dataset to determine if there was a statistically significant difference in the distribution of the scoring metric between the TP and TN peptidase-inhibitor pairs. The scoring metrics for each predicted peptidase-inhibitor structure was used to conduct a receiver operating characteristic (ROC) area under the curve (AUC) analysis to assess the effectiveness of a scoring metric from the predicted complexes in distinguishing between the TP and TN peptidase-inhibitor pairs in each dataset. In each iteration, a ROC curve was generated, and the AUC was calculated using 4/5-fold of the data to evaluate classification performance; an optimal threshold for classification was determined using Youden’s J statistic, maximizing sensitivity and specificity. This threshold was applied to classify the remaining 1/5-fold of the data, which was compared to the TP and TN labels. The input for the ROC-AUC analysis were the scores for a scoring metric from each peptidase-inhibitor pair in a test data set. The output from the five-fold cross validation ROC-AUC analysis for each input included the average optimal threshold for the scoring metrics, the average AUC, the average true positive rate (TPR) and false positive rate (FPR) at the optimal thresholds, and the average classification accuracy of the 1/5-fold using the optimal thresholds. These output parameters were recorded for each of the six scoring metrics (i.e., mean and max ipTM, LIS and LIA) for each test dataset.

### Peptide-library preparation

The peptide library was built by *in silico* digestion of the *Capaea nemoralis* proteome with trypsin, subtilisin, thermolysin, and pepsin (at pH 1) using the algorithm “rapid peptides generator” (RPG) ^33^ followed by prediction of antifungal potential ^34^. From this prediction, the top 100 candidates were selected for antifungal testing.

### Predicting inhibitor candidates for C. neoformans peptidases

Using our computational pipeline, we predicted the interaction scores between the *C. nemoralis* peptide library and three virulence-related peptidases in *C. neoformans*: Rim13, May1, and CnMpr1. Based on classification as TP, two predicted peptidase inhibitors were prioritized for Rim13 and May1 and four predicted peptidase inhibitors were prioritized for CnMPr1. The peptidase inhibitors were synthesized by PepMic (China) and the Structural and Biophysical Core Facility from The Hospital for Sick Children (Canada) with a purity >90%; confirmed via HPLC (Fig. S1 and S2). For HPLC purity assessment, a SHIMADZU Inertsil ODS-SP (4.6*250mm*5um) column was used to separate the peptides with Buffer A (0.1% TFA in 100% water) and Buffer B (0.1% TFA in 100% acetonitrile) across a 60 min gradient at 1 mL/min with OD_220nm_ measured. Peptides were dissolved in 30% Acetonitrile (ACN) in water combined with 0.1% trifluoroacetic acid (TFA) to a concentration of 10 mg/mL. From this stock, aliquots at specified concentrations were prepared and stored at −20 °C.

### Media, strains and growth conditions

*C. neoformans* variety *grubii* strain H99 (Wild Type) was used for all cultures related to virulence factor assays. The lab-evolved (LE) fluconazole-resistant (64 μg/mL) strain was used for resistance assays ^35^. H99 was routinely stored on yeast peptone dextrose (YPD) agar (1% yeast extract, 2% bacto-peptone, 2% D-glucose, 2% agar) and stored at 4 °C. The LE strain was maintained on YPD plates supplemented with 64 μg/mL fluconazole and 34 μg/mL chloramphenicol. From these plates, one colony was resuspended in YPD broth and grown overnight at 37 °C and 200 rpm. Overnight cultures of *C. neoformans* in YPD were washed and resuspended in Yeast Nitrogen Base (YNB, 0.67% of yeast nitrogen base, 0.5 % of glucose) (Sigma Aldrich) to a concentration of 10^5^ cells/mL and used for subsequent analyses.

### Gene deletion and complementation

Coding DNA sequences (CDS) for each peptidase were identified using FungiDB ^36^ with genetic deletion strains generated using biolistic transformation and double-joint PCR methodology ^37^. Briefly, upstream and downstream homologous regions (approximately 1000 bp) of CNAG_05601 (Rim13), CNAG_05872 (May1), and CNAG_04735 (CnMpr1) and were joined via double-joint PCR to the nourseothricin (NAT) resistance cassette (generously provided by Dr. J.P. Xu, McMaster University) ^38^. Primers and plasmids provided (Table S1). DNA construct was inserted in the genome via biolistic transformation ^39^. Stable transformants were selected on YPD-NAT (100 µg/mL) agar and confirmed by diagnostic PCR, and two independent mutants were verified for correct genomic deletion by Southern blot.

Genetic complementation was performed using plasmids generated by Gibson assembly (New England Biolabs): gene promoter (approximately 1000 bp upstream of start codon), coding region, 4X-FLAG tag added to the 3’ end of the open reading frame, and Hygromycin B (HYG) resistance cassette, and 3’ homologous region into pSDMA58 (generously provided by Dr. J. Fraser, University of Queensland). Plasmids were inserted by electroporation into both mutant and WT strains. Briefly, overnight cultures were resuspended 1:50 into 100 mL YPD and incubated for 6 h at 30 °C and 200 rpm to OD_600nm_ = 0.6-1.0. Cells were collected by centrifugation, washed, resuspended in cold elution buffer (EB) (10 mM Tris-HCl (pH 7.5), 1 mM MgCl_2_, 270 mM Sucrose) supplemented with 1 mM dithiothreitol (DTT) and incubated on ice for 30 min followed by washing and resuspension in 300 µL of EB buffer. Five microliters of DNA was combined with 45 µL of fungal cells in sterile 1.5 mL tube and placed on ice. Cells and DNA were transferred to an ice cold electroporator cuvette and electroporated with a BioRad gene pulser at 0.45 kV, 125 uF, 400-600 Ohms. Then, 1 mL YPD broth supplemented with 0.5 M sorbitol was transferred into the cuvette and left to recover for 90 min at 30 °C. Cells were plated and stable transformants were selected on YPD-HYG (200 µg/mL) agar, confirmed by PCR.

### Expression and purification of peptidases

Expression of peptidases was achieved by mimicking natural conditions for each enzyme. Briefly, each strain containing the tagged enzyme was grown in 5 mL of YPD broth overnight at 30 °C, 200 rpm. Cells were sub-cultured to 250 mL YPD broth overnight at 30 °C and 200 rpm, washed by gentle centrifugation (1000 x g for 5 min) and resuspended in 250 mL of expression media for 24 h at 30 °C and 200 rpm as follows: Rim13 (CNAG_05601) in YNB pH 7.4; May1 (CNAG_05872) and CnMpr1 (CNAG_04735) were expressed in YNB pH 5 supplemented with 0.1% of Bovine Serum Albumin (BSA). All media contained 200 µg/mL HYG. Cells were collected by centrifugation and resuspended in 25 mL of cold tris-buffer saline (TBS) supplemented with 5 mM of phenylmethylsulfonyl fluoride (PMSF) and aliquoted in high impact 1.5 mL tubes. One scoop (approximately 1 g) of glass beads was added to each 1.5 mL tube before lysis using a bullet blender at 1200 rpm for 5 min at 4 °C. Cells were collected by centrifugation at 15000 x g for 10 min at 4 °C. The supernatants were collected and filtered using 0.45 µm filter-membranes and stored at −20 °C until purification. Protein production was confirmed by Western Blot using an anti-FLAG antibody (Sigma, USA).

To purify the enzymes, 100 µL of anti-FLAG resin was washed and equilibrated with TBS pH 7.4 before mixing with 10 mL of supernatant. Anti-FLAG resin and samples were incubated for 24 h with rotation at 4 °C. After this time, the resin was precipitated with gentle centrifugation (200 x g for 2 min), and supernatant was labeled as ‘flow-through’. The resin was washed once with TBS before elution with acid glycine at pH 3 for 5 min with rotation. Elution steps were repeated twice, with each elution resuspended in 10% (v/v) of 1 M NaOH to increase the pH and avoid denaturation. The presence of each protein was confirmed by Western blot and concentrated using a 10 kDa cutoff concentrator column and resuspended in PBS.

### Enzymatic assays

Inhibitory activity was assessed by incubating each candidate inhibitor at different concentrations with the peptidases for 10 min in the corresponding buffer across temperatures (Table S2). Enzymatic activity was measured by adding the corresponding substrate using manufacturer instructions and monitoring the product’s appearance over time (every 5 min) until linearity was lost (approximately one hour) by fluorescence measurements using a plate reader (Synergy-H1, Biotek). Excitation and emission wavelengths and reagents are reported (Table S2). Inhibitory Concentration of 50% (IC_50_) values were obtained by interpolation of data corresponding to the lineal region of the dose-response curve.

### Fungal growth and thermotolerance assays

To assess the effect of peptidase inhibitor candidates on *C. neoformans* growth at 37 °C, 6 μL of extract serial dilution series was mixed with 194 μL of fungal cells (previously resuspended in YNB) in a 96-well plate. Growth curves were measured using a plate reader (Synergy-H1, Biotek) at 200 rpm for 72 h, followed by optical density (OD_600nm_) readings every 30 min. All experiments were performed in three biological and two technical replicates.

### Polysaccharide capsule induction

Overnight cultures of *C. neoformans* H99 from YPD were sub-cultured in YNB and incubated overnight at 37 °C with 200 rpm shaking. Production of polysaccharide capsule was induced as previously described ^22^. Briefly, peptide inhibitors and 10^7^ cells/mL of *C. neoformans* were cultured in 5 mL of low iron media (LIM; 0.5% L-asparagine, 0.4% HEPES, 0.04% K_2_HPO_4_, 0.008% MgSO_4_·7H_2_O, 0.2% NaHCO_3_, and 0.025% CaCl_2_·2H_2_O) and incubated for 72 h at 37 °C. Visualization of cells was performed by mixing cells with India ink using a 1:1 ratio on microscope slides and visualizing with a Differential Interference Contrast (DIC) microscope and a 63X oil objective. Capsule production was quantified using a ratio of total cell size (with capsule) to cell size (without capsule). All measurements were obtained using three biological replicates with 40-60 cells assessed per condition.

### Blood-Brain Barrier integrity and crossing assays

Crossing of the BBB was tested using a previously developed model ^40,41^ with slight modifications. Briefly, Induced Pluripotent Stem Cell (IPSC) Line #41658 was obtained from the NINDS repository (ND 41658, male), defrosted and plated in a six-well plate. Once the cells reached 70-80% confluence, the Brain Microvascular Endothelial Cell (BMEC) differentiation protocol was initiated ^42^. Cells were subjected to the detachment solution (accutase) for three minutes, centrifuged, resuspended and counted to be seeded in a six-well plate at 11,000 cells/cm^2^. After 24 h, cells were switched into E6 media (TeSR™-E6, STEMCELL Technologies); E6 media was changed every 24 h until the fifth day when the media was changed to EC (Human endothelial serum free medium, 1% platelet-poor plasma derived bovine serum, 20 ng/mL human basic fibroblast growth factor (bFGF) and 10 µM retinoic acid). On the sixth day, transwells were coated with collagen IV (1:10) and fibronectin (1:20) attachment substrates and on the seventh day, cells were detached from the plate by incubation with accutase for 25 min, centrifuged, resuspended in EC++ media (EC media supplemented with 3% 70 kDa dextran, 20 µM phosphodiesterase inhibitor, 400 µM Dibutyryl cyclic AMP, 10 µM retinoic acid), and 10 µM rock inhibitor γ-27632) and seeded in the transwell at 195,000 cells/well.

Cryptococcal cells were sub-cultured in YPD at 37 °C overnight at 200 rpm, washed by centrifugation with PBS and resuspended in EC+ media to a final concentration of 1.56 × 10^6^ cells/mL. From this stock, 250 µL of cells were added to each insert to 3.9 × 10^5^ cryptococcal cells or a multiplicity of infection (MOI) of 2:1. Each insert was loaded in a 24 well plate containing 700 µL of EC media. Peptide inhibitors were added, and inserts were incubated for 24 h at 37 °C in 5% CO_2_. Then, 100 µL was collected from the bottom of the well, serially diluted and plated on YPD for 48 h at 30 °C for colony forming unit (CFU) counting.

### Fluconazole resistance

Cryptococcal cells were prepared as outlined above for growth assays and the experiment was performed using a checkerboard assay ^43^. Briefly, cells from an overnight culture of the LE strain were collected, washed and resuspended in YNB media to a final concentration of 10^6^ cells/mL. Each peptidase inhibitor candidate (from 32 to 0 µg/mL) was mixed with fluconazole (from 128 to 0 µg/mL) in the presence of the LE strain in 96-well plates, wrapped in aluminum foil, and incubated at 30 °C and 800 rpm for 48 h. Growth was discontinuously followed at OD_600nm_ every 24 h. Optical density values for each condition were normalized against the corresponding control (i.e., cells without treatment). Synergy between peptide inhibitors and fluconazole was assessed by the fractional inhibitory concentration index (FICI) model ^44^.

### Macrophage fungal clearance model

To assess the susceptibility of the WT (H99) strains against immortalized macrophages derived from BALB/c mice in the presence of the peptidase inhibitors, the fungal cells were grown as previously described with minor variations ^45^. Briefly, macrophages were normalized to 50,000 cells/mL using Dulbecco’s Modified Eagle Medium (DMEM) supplemented with 10% fetal bovine serum (FBS), penicillin and streptomycin and statically incubated at 37 °C, CO_2_ 5% for 48 h. The *C. neoformans* strains were cultured in liquid YPD at 37 °C and 200 rpm for 18 h and sub-cultured overnight in the same conditions. Cryptococcal cells were collected by centrifugation and washed two times with PBS. Using a hemocytometer, cells were normalized to 1 × 10^6^ cells/mL in 1 mL of DMEM. Cryptococcal cells were opsonized by adding 1 µg of anti-glucuronoxylomannan (GXM) antibody (1 µg/mL) ^46^ and incubated for one hour at 37 °C and 5% CO_2_.

For infection, macrophages were gently washed with 1 mL of PBS and diluted in DMEM with *C. neoformans* (10^6^ cells/mL). After static incubation for 90 min at 37 °C and 5% CO_2_, cells were washed two times with PBS and diluted in 1 mL DMEM with each inhibitor and incubated for 18 h at 37 °C and 5% CO_2_. Macrophages were washed twice with PBS and incubated with 1.2% Triton X-100 for 10 min at room temperature. From each well, 1 mL was collected and serially diluted in PBS using a 10-fold series until 10 cells/mL was achieved. From each dilution, 100 µL was plated on YPD-agar plates, allowed to dry at room temperature for 10 min, and incubated at 30 °C for 48 h. CFU were counted on each condition and dilution. Experiments were performed in biological triplicate and technical duplicate.

### Cytotoxicity assay

To analyze the cytotoxic effect of the inhibitors against mammalian cells, we assessed the lactate dehydrogenase (LDH) activity of BALB/c macrophages as previously described ^45^. Peptide inhibitor candidates at 10 µg/mL were combined with 1 mL DMEM-containing macrophages using 24-well plates and, statically incubated at 37 °C, 5% CO_2_ for 18 h. Triton (1.2%) was added to the non-treated wells for total death and incubated at room temperature for 30 min. LDH substrate (NAD^+^) (Sigma-Aldrich, US) and samples were mixed using a 1:1 ratio (*v:v*) in 96-well plates and incubated at room temperature for 20 min before adding a stopping solution (Sigma-Aldrich, US). LDH activity was quantified by measuring OD_450nm_. Media only and Triton 1.2% were used as blanks for extract-containing wells and total death replicates. Each experiment was performed using three biological and two technical replicates.

### Biofilm formation and quantification

Biofilm formation was carried out following the protocol previously described, with minor modifications ^47^. Overnight cultures of *C. neoformans* were washed twice with PBS and resuspended in DMEM, supplemented with glutamine and sodium pyruvate, to an OD_600nm_ of 1.0 (approximately 1 × 10^7^ cells /ml). A 96-well plate was prepared by adding 100 µl of a corresponding 1:25 peptidase inhibitor candidate dilution in DMEM to each well, followed by 100 µl of the adjusted cell suspension. The plate was incubated statically at 37 °C for 48 h to allow biofilm formation. After incubation, the supernatant was carefully removed, and wells were washed three times with 200 µl of PBS. Plates were air-dried at room temperature followed by addition of 100 µl of 0.2 % (w/v) crystal violet solution to each well for staining and incubated for 10 min at room temperature. Excess of crystal violet was removed by washing the wells three times with deionized water, and the plates were dried at room temperature. To quantify biofilm biomass, 200 µL of 30 % (v/v) acetic acid was added to each well to solubilize the crystal violet. After a 5-minute incubation, the supernatant was transferred to a new plate, and absorbance was measured at 550 nm. Three biological and two technical replicates were included for this experiment.

### Galleria mellonella larvae in vivo model

*Galleria mellonella* (i.e., greater wax moth) larvae infection model for *C. neoformans* was performed as previously described with some modifications ^48^. Briefly, upon arrival, larvae were washed by five quick submersions in bleach 0.1% and ten times in water. After the larvae dried, those in good condition (i.e., yellow color and high activity) were placed in petri dishes (approximately 20 each) and incubated at 37 °C overnight. The next day, overnight cultures of *C. neoformans* were washed with PBS twice and normalized to 10^6^ cells/mL. Then, larvae were inoculated with 10 µL of culture (10^4^ cells) using a Hamilton syringe and incubated at 37 °C. If applicable, larvae were allowed to rest for 4 h before applying 10 µL of peptidase inhibitor candidates with or without fluconazole (10 mg/kg; diluted in PBS) at a final concentration of 10 mg/kg. Every 24 h, dead larvae (i.e., black in color or no movement after stimulation) or pupa were counted and separated from the rest. Pupas were not considered for analysis. Survival curves were built using the Kaplan-Meier estimator.

### Mass spectrometry analysis

Profiling of *C. neoformans* cells was performed as previously described for total proteome samples ^49^. Briefly, cells were collected by centrifugation, washed with PBS and lysed using a sonication. Proteins were precipitated overnight using 80% acetone and 25 µg of proteins were enzymatically digested using a trypsin/Lys-C mixture, followed by desalting and purification using STop And Go Extraction (STAGE)-tips ^50^. Peptides were prepared as we previously described with the Pierce^TM^ fluorometric peptide quantification assay (Thermo Fisher Scientific), normalized to 35 ng/µL, and Indexed retention time (iRT) synthetic peptides (Biognosys AG) were spiked into each sample at 1:5 (iRT:sample) prior to LC-MS/MS analysis ^51^.

Peptides were separated using the Thermo Scientific™ Vanquish™ Neo™ UHPLC system in a trap-and-elute injection configuration at 60 samples per day (23.35 min gradient) and analyzed with the Thermo Scientific™ Orbitrap™ Astral™ Zoom mass spectrometer. A Thermo Scientific™ EASY-Spray™ HPLC ES906PN column (150 μm x 15 cm, 2 μm pore size) was used for analytical separation. Mobile phases: 0.1% formic acid in water (A) and 0.1% formic acid in 80% acetonitrile (B). Eluted peptides were analyzed using narrow-window data-independent acquisition (nDIA) methodology on the Orbitrap™ Astral™ Zoom mass spectrometer. Parameters for full scan MS1 and nDIA MS2, along with gas-phase fractionation injections of a *C. neoformans* reference culture were set as we previously reported ^51^.

### Mass spectrometry data processing

All mass spectrometer output files were analyzed using Spectronaut v19.8.250311.62635 (Biognosys AG). All mass spectrometer output files were analyzed using directDIA in Spectronaut v19.8.250311.62635 (Biognosys AG) with the protein FASTA files against *C. neoformans* var. *grubii* serotype A (strain H99/ATCC 208821; UniProt UP000010091; 7,429 sequences). Default search parameters were applied during the directDIA analysis search: trypsin enzyme specificity with a maximum of two missed cleavages, carbamidomethylation of cysteines (fixed modification), and oxidation of methionine and N-acetylation of proteins (variable modifications). Precursor ion mass and fragment ion tolerance values were set as the Dynamic default within Spectronaut. Spectral matching of the peptides was performed with a false discovery rate (FDR) of 1% identified proteins with a minimum of two peptides for protein identification. Quantification was based on MS2 peak areas, no imputation strategy was used, and local normalization was applied.

### Mass spectrometry data analysis and visualization

Statistical analysis and data visualization of the proteomics data were performed using Perseus (version 2.1.4.0) and ProteoPlotter ^52,53^. Data was prepared by filtering for reverse database matches, contaminants, and proteins only identified by site, followed by log_2_ transformation of protein intensities. Filtering for valid values (three of four replicates in at least one group) was performed, missing values were imputed from the normal distribution (width, 0.3; downshift, 1.8 standard deviations), and group values were averaged. Significant differences were evaluated by a Student’s *t*-test (p-value ≤ 0.05) with multiple-hypothesis testing correction using the Benjamini-Hochberg (FDR = 0.05) with S_0_ = 1 (2-fold difference between comparisons) ^54^. Proteomics profiling was performed in biological quadruplicate.

### Statistics

For phenotypic and *in vitro* assays (i.e., growth, capsule, and biofilm), data were visualized and statistically analyzed using GraphPad Prism version 9.0 (GraphPad Software, Inc., USA; https://www.graphpad.com/). P values of ≤ 0.05 were considered significant.

## Results

### Scoring metric distributions classify putative inhibitors as true positive or true negative

To design putative inhibitors of the target cryptococcal peptidases, we developed a curated database of 108 known peptidase-inhibitor complexes as a testing dataset to process through our computational pipeline. The pipeline integrated AFM^24,25^ with Alpha-Pulldown^26^ and AlphaFold2^27^ for paralleled multichain protein complex prediction to model enzyme-inhibitor interactions (Fig. S3). The peptidase (i.e., cysteine, aspartic, metallo) and inhibitor amino acid sequences were used as input. Inhibitors were divided into two classes: peptides (< 30 amino acids) and proteins (>30 amino acids). Scoring metrics for the mean interface predicted template modeling (ipTM), localized interaction score (LIS), and localized interaction area (LIA) scores were evaluated. For ipTM, which measures the accuracy of the predicted relative positions of the subunits forming the protein-protein complex, values higher than 0.8 represent confident high-quality predictions (i.e., true positives [TP]), while values below 0.6 suggest likely a failed prediction (i.e., true negatives [TN]); values between these scores are indistinguishable as TP or TN. Additionally, LIA and LIS were used to filter the prediction solutions by consideration of the active site information of the selected protease-inhibitor complexes. For the peptide inhibitor test data set (<30 amino acids), the ipTM, LIS, and LIA scores for the TP group were significantly higher than the TN group (Fig. 1A-C). Similarly, for the protein inhibitor test data set (>30 amino acids) the scores for the TP group were significantly higher than the TN group across all evaluated scoring metrics (Fig. 1D-F). Notably, the overlap of scores between TP and TN was higher for peptide vs. protein inhibitors, suggesting a higher false positive rate (FPR) and false negative rate (FNR) for peptide-based inhibitors. Based on these results, we determined that AFM differentiates between TP and TN peptidase-peptide or -protein inhibitor complexes using the mean ipTM, LIS, and LIA scores.

**Figure 1:**
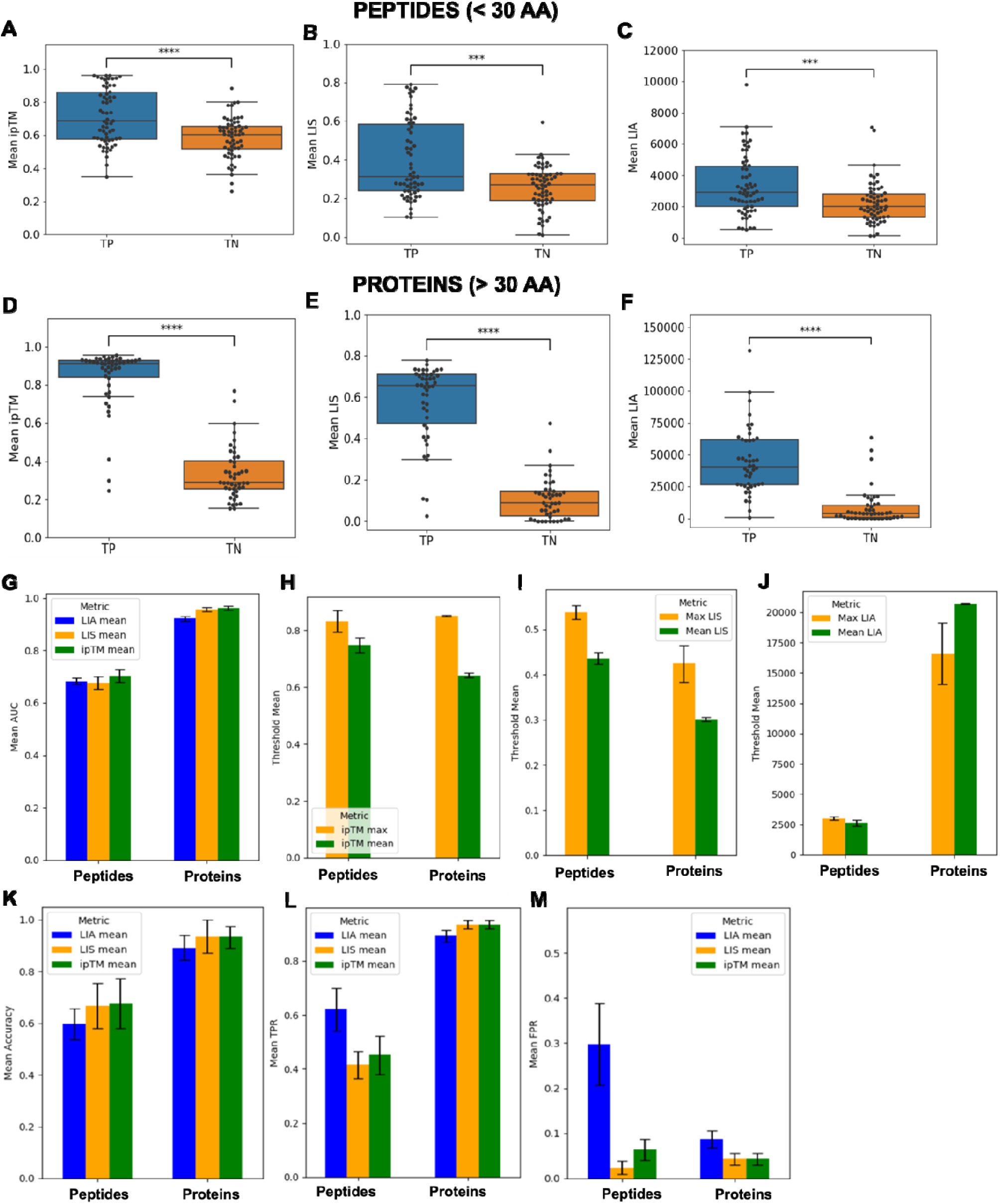
Computational pipeline scoring metric extraction and evaluation and AUC and accuracy measurements between TP and TN for peptide and protein inhibitors. **A)** Mean ipTM for each peptidase-peptide pair in each of the TP and TN groups. **B)** Mean LIS for each peptidase-peptide pair in each of the TP and TN groups. **C)** Mean LIA for each peptidase-peptide pair in each of the TP and TN groups. **D)** Mean ipTM score for each peptidase-protein score in each of the TP and TN groups. **E)** Mean LIS score for each peptidase-protein score in each of the TP and TN groups. **F)** Mean LIA score for each peptidase-protein pair in each of the TP and TN groups. **G)** Mean AUC for the scoring metrics for each of the evaluation data sets. **H)** Mean optimal threshold for differentiating TP and TN peptidase-inhibitor pairs calculated from the mean and max ipTM. **I)** Mean optimal threshold for differentiating TP and TN peptidase-inhibitor pairs calculated from the mean and max LIS. **J)** Mean optimal threshold for differentiating TP and TN peptidase-inhibitor pairs calculated from the mean and max LIA. **K)** Mean classification accuracy in the hold-out fold using the optimal threshold calculated for the mean scoring metrics. **L)** Mean TPR from ROC-AUC analysis at the optimal threshold. **M)** Mean FPR from ROC-AUC analysis at the optimal threshold. Peptide data set consisted of 62 TP and 62 TN. Protein data set consisted of 46 TP and 46 TN. Error bars indicate standard deviations. The significance of the group difference from a two-sided Mann-Whitney U-test for each scoring metric is denoted by the asterisk (*** = p <0.0001, *** = p <0.001, ** = p <0.01, * = p<0.05, ‘NS’ = p>0.05). Figures were prepared in R.

### Area under the curve and accuracy measurements differentiate clustering of TP from TN for peptide and protein inhibitors

The mean area under the curve (AUC) was calculated based on the mean ipTM, LIS, and LIA scoring metrics from the predicted peptidase-inhibitor complexes in each test set to assess the overall classification performance for each score derived from the predicted complex. We observed a higher mean AUC across each scoring metric from the protein inhibitor test data set, with mean AUC values above 0.9 compared to the peptide inhibitor test set where the mean AUC values were between ∼0.65 – 0.7 (Fig. 1G). These data suggest that AFM performed best at differentiating TP and TN peptidase-inhibitor pairs from the protein inhibitor data set compared to the peptide inhibitor data set. Next, we performed a comparison between the mean optimal threshold value for the max and mean ipTM, LIS, and LIA for the peptide and protein inhibitor test sets to assess the viability of these thresholds to differentiate unclassified TP and TN peptidase-inhibitor pairs. We observed that the average optimal scoring metric (i.e., threshold) for the max ipTM and max LIS were higher than the mean values for peptide and proteins inhibitors with relatively low standard deviation (Fig. 1H,1I). The average optimal threshold for the max and mean LIA scores were comparable for peptides but max values were lower than mean for protein inhibitors (Fig. 1J). The low standard deviations in the mean optimal scores for the ipTM, LIS, and LIA scores in the peptide and protein inhibitor test sets suggest a convergence of an optimal threshold that could be used to differentiate unclassified TP and TN peptidase-inhibitor pairs. Notably, however, the higher standard deviation in the mean threshold for the max LIA suggests less convergence in a single threshold, potentially indicating less reliability in using this score to differentiate TP and TN peptidase-protein inhibitors.

To evaluate classification accuracy using the optimal threshold of the scoring metrics, we quantified classification accuracy of the hold-out fold (i.e., 1/5-fold during each iteration of cross validation). With accuracy approaching 90%, we observed AFM to provide clearer differentiation between TP from TN for protein inhibitors compared to peptide inhibitors (accuracy around 60%) based on mean scores (Fig. 1K) with very similar results observed for the max scores (Fig. S4). Notably, for both the peptide and protein inhibitor test sets, the three-scoring metrics showed similar mean accuracies compared to each other with a slight reduction for LIA mean. Next, we evaluated the mean true positive rate (TPR) at the optimal thresholds for each of the scoring metrics. Like the mean classification accuracy in the hold-out fold, AFM showed higher TPR for the protein inhibitor set (approx. 0.9) compared to the peptide inhibitor set (between. 0.4-0.63) (Fig. 1L). These results indicate that across different subsets of the protein inhibitor test set, it is possible to correctly classify most TP peptidase-inhibitor pairs using the three scoring metrics while approximately only half of the TP peptidase-inhibitor pairs from the peptide data set were successfully classified. When assessing the associated mean FPR at the optimal score for each scoring metric, each of the mean ipTM, LIS, and LIA had a mean FPR below 0.1 in the protein inhibitor test set (Fig. 1M). This indicates that for peptidase-inhibitor pairs in the protein inhibitor data set, under 10% of the TN pairs were miscategorized as TP using the scoring metrics. For the peptide inhibitor test set, we observed that the mean ipTM and LIS scores had a mean FPR below 0.1; however, the mean LIA had a higher mean FPR (approx. 0.3), suggesting almost a 30% misclassification of TNs as TPs based on the mean LIA score for peptidase-peptide inhibitors. These data show that application of the three mean scoring metrics when screening protein inhibitors minimize the FPR below 10% while for peptide inhibitors, the mean ipTM and LIS scores should be used to minimize the FPR below 10%.

### Structure-guided prediction of putative inhibitors toward cryptococcal peptidases prioritizes candidates for experimental evaluation

Given the global health implications of fungal diseases and the limited arsenal of antifungal agents available, we applied our computational workflow using optimal thresholds for the mean ipTM and LIS scores to predict and screen 108 putative inhibitors toward three virulence-related peptidases in *C. neoformans:* Rim13, May 1, and CnMpr1. For Rim13, with known roles in capsule production, iPTM and LIS scores higher than the proposed threshold (Table S3), prioritized two putative inhibitors: P2221 and CPI-1. Based on pipeline structure prediction, P2221 binds to the active site region of Rim13 characterized by the presence of its three catalytic residues: C185, H354 and D374 (Fig. 2A). Here, cysteine 8 (C8) from P2221 is predicted to form a disulfide bridge with the catalytic C185 as a warhead while the remaining chain interacts with other amino acids, such as V182 and A377, of the exposed region. However, based on this prediction, amino acids located at the N- and C-termini of P2221 appeared remote from the active site with no clear role in enzyme binding or inhibition. Given that Rim13 is an intracellular target, requiring an inhibitor to cross the fungal cell wall and membrane to act on the enzyme, we designed a shorter version of P2221 with similar binding strength but improved pharmacological properties by performing single amino acid substitutions along the peptide backbone and shortening the peptide length to promote optimized interactions (e.g., higher LIS) within the binding site. Using this strategy, we substituted isoleucine 10 for methionine (I10M6) and moved C8 to the peptide center (C8 to C4); three amino acids upstream and downstream of the peptide center were unchanged (Fig. S4A; Table S4). The modified peptide with optimized interaction and size was deemed CPI-1. Like P2221, CPI-1 is predicted to bind in the active site region of Rim13 with C4 having an important role due to its interaction with the catalytic C185 (Fig. 2B). Based on these properties and modifications, we predict that P2221 and CPI-1 will target and inhibit Rim13 to reduce enzyme activity and impair capsule production.

**Figure 2:**
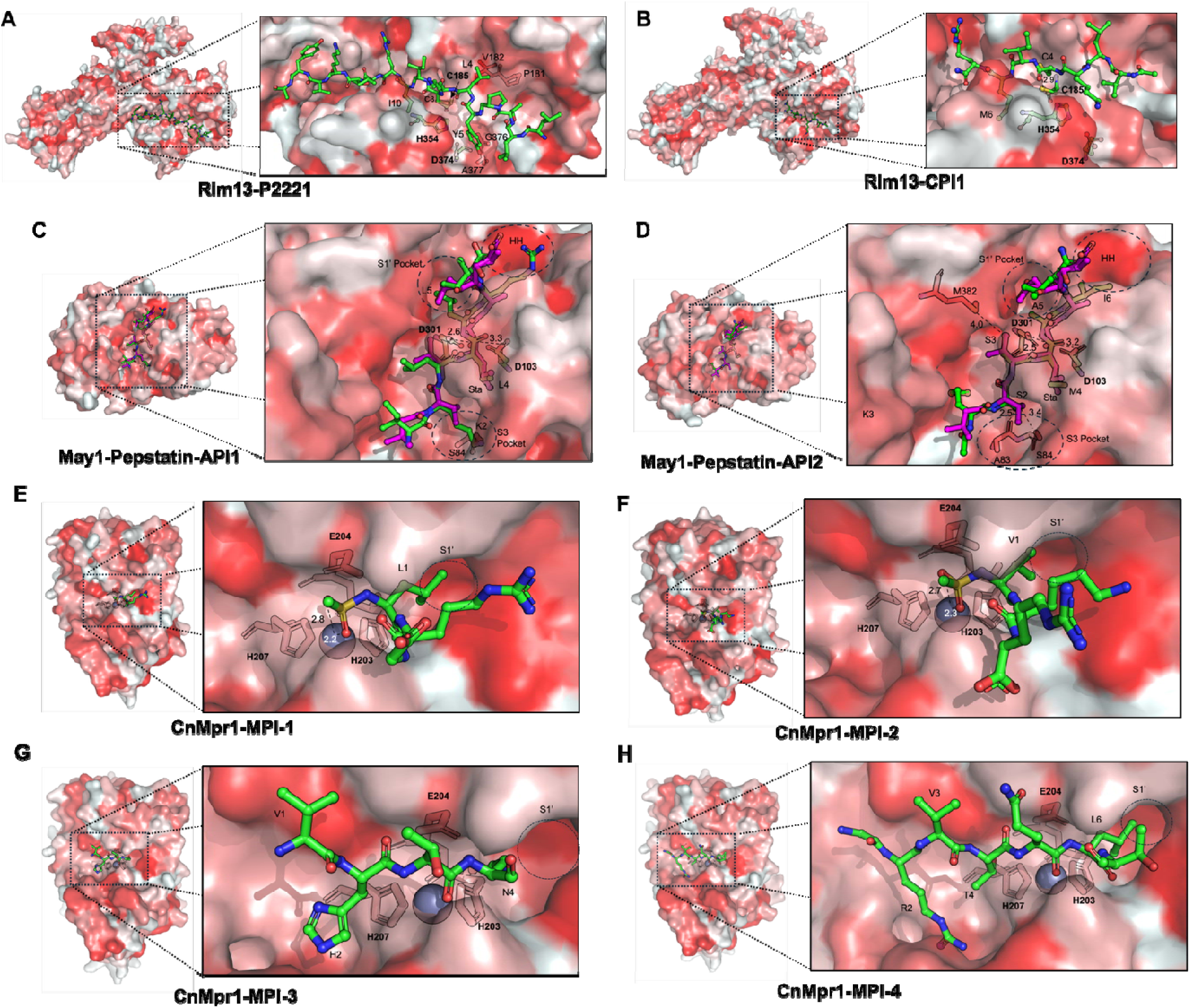
Predicted putative inhibitors for key cryptococcal peptidases. **A** and **B**) Predicted binding of P2221 and CPI-1 with Rim13. **C** and **D**) Superimposition of the predicted binding between A I-1 and API-2 with May1, respectively. Complex between PEP-A and May1 was derived from PDB: 6R6A. PEP-A is colored magenta. **E-H)** Prediction of binding between MPI-1, 2, 3 and 4 with CnMpr1, respectively. Ligands are shown as sticks and colored by element: carbons are green, oxygens are red, nitrogens are blue, and sulphurs are orange. Members of the active site are highlighted in bold font. Amino acids were numbered from N- to C-termini. Distances are shown in Angstroms. Protein images were prepared with Pymol (https://pymol.org/).

Next, for May1, with known roles in pH response and macrophage survival, iPTM and LIS interaction scores (Table S5) prioritized trypsin 3075 and trypsin 735 for optimization using single amino acid substitutions and rational design for two inhibitors, API-1 and API-2 (Fig. S4B, S4C; Table S6, S7). Given that Pepstatin A is a known inhibitor of aspartic peptidases, including May1, we superimposed May1-API-1 complex with the complex May1-Pepstatin (PDB: 6R6A) to assess similitudes and predict the P1 position for substitution with statine. We observed that API-1 has a similar binding position as Pepstatin-A in the active site of May1 (Fig. 2C). Notably, leucine 4 (L4) from API-1 is in same position of statine (from Pepstatin A), with close proximity to the catalytic dyad, D301 and D103. Furthermore, leucine 5 (L5) occupies the S1’ pocket and lysine 2 (K2) interacts with serine 84 (S84) within the S3 region of May1 via hydrogen bonds. API-2 is predicted to dock similarly to API-1 with a methionine 4 (M4) analog to statine close to the catalytic dyad (Fig. 2D). Moreover, serine 2 (S2) is predicted to occupy the S3 pocket but has an Ala residue proximal to but not long enough to interact with the S1’ pocket. Likewise, isoleucine 6 (I6) is oriented towards the high hydrophobic area located at the edge of the active site. Based on these properties and modifications, we predict that API-1 and API-2 will target and inhibit May1 to reduce enzyme activity and survival of *C. neoformans* within macrophages.

For CnMpr1, with known roles in dissemination and BBB crossing, ipTM and LIS interaction scores prioritized two candidate inhibitors (Table S8). Specifically, we substituted the N-termini for sulfonamide, which have known metallopeptidase inhibitory activity^55^, to create MPI-1 and MPI-2 (Fig. S5A). In parallel, we performed a single amino acid substitution analysis to improve the binding between the top predicted inhibitors and CnMpr1 to create MPI-3 and MPI-4 (Fig. S5B; Table S9). MPI-1 and MPI-2 are peptidomimetic with a sulfonamide group in the N-termini, which is predicted to bind to Zn within the CnMpr1 active site (Fig. 2E, 2F). In both cases, MPI-1 and MPI-2 possess an aliphatic residue (Leu and Val, respectively) oriented towards the S1’ pocket. On the other hand, MPI-3 is a tetrapeptide that is predicted to fold around the Zn^2+^ ion at the active site (Fig. 2G). MPI-3 also possesses a Val residue in position 1 (V1) that is predicted to interact with a hydrophobic patch close to the active site. Notably, there are no residues oriented toward the S1’ pocket in MPI-3. Similarly, MPI-4 is a heptapeptide that is predicted to fold around the Zn^2+^ ion but possesses a Leu in position 6 oriented towards the S1’ pocket (Fig. 2H). MPI-4 also has a Thr residue (T4) in position 4 close to the Zn^2+^, and a Val group (V3) predicted to interact with the hydrophobic patch above the active site. Based on these properties and modifications, we predict that MPI-1, MPI-2, MPI-3, and MPI-4 will target and inhibit CnMpr1 to reduce enzyme activity and impair BBB crossing. Together, our computational pipeline designed putative inhibitors based on the ipTM and LIS scoring metrics for each of the targeted cryptococcal peptidases with modifications to increase target specificity and inhibitory activity.

### Rim13 inhibitors reduce the ratio of polysaccharide capsule to cell size

To evaluate the efficacy of our computationally designed peptidase inhibitors through modulation of Rim13 activity, we focused on the role of the target peptidase in polysaccharide capsule production. Specifically, Rim13 is a C2 cysteine peptidase involved in the activation of Rim101, which regulates integrity of the cell wall and capsule formation ^30^. To set a baseline of phenotypic disruption, we generated a gene deletion strain of *rim13* (CNAG_05601) in the *C. neoformans* H99 background to confirm its role in capsule production (Fig. S6A, S7A). As anticipated, we observed a significant reduction in capsule, cell body size, and the ratio of capsule to cell size for *rim13*Δ compared to WT (Fig. 3A). Next, we expressed, purified, and confirmed production of FLAG-tagged Rim13 (Fig. S8A) and performed inhibition assays of the peptidase in the presence of the putative inhibitors. We observed that both P2221 and CPI-1 inhibited Rim13 with inhibitory concentration at 50% (IC_50_) of 2.5 and 5.5 µM, respectively (Fig. 3B). Additionally, we tested E64 (a known cysteine peptidase inhibitor with capsule disruption properties^23^) and observed an IC_50_ of 1.1 µM against Rim13, confirming the cysteine-type enzymatic activity of the prioritized inhibitors.

**Figure 3:**
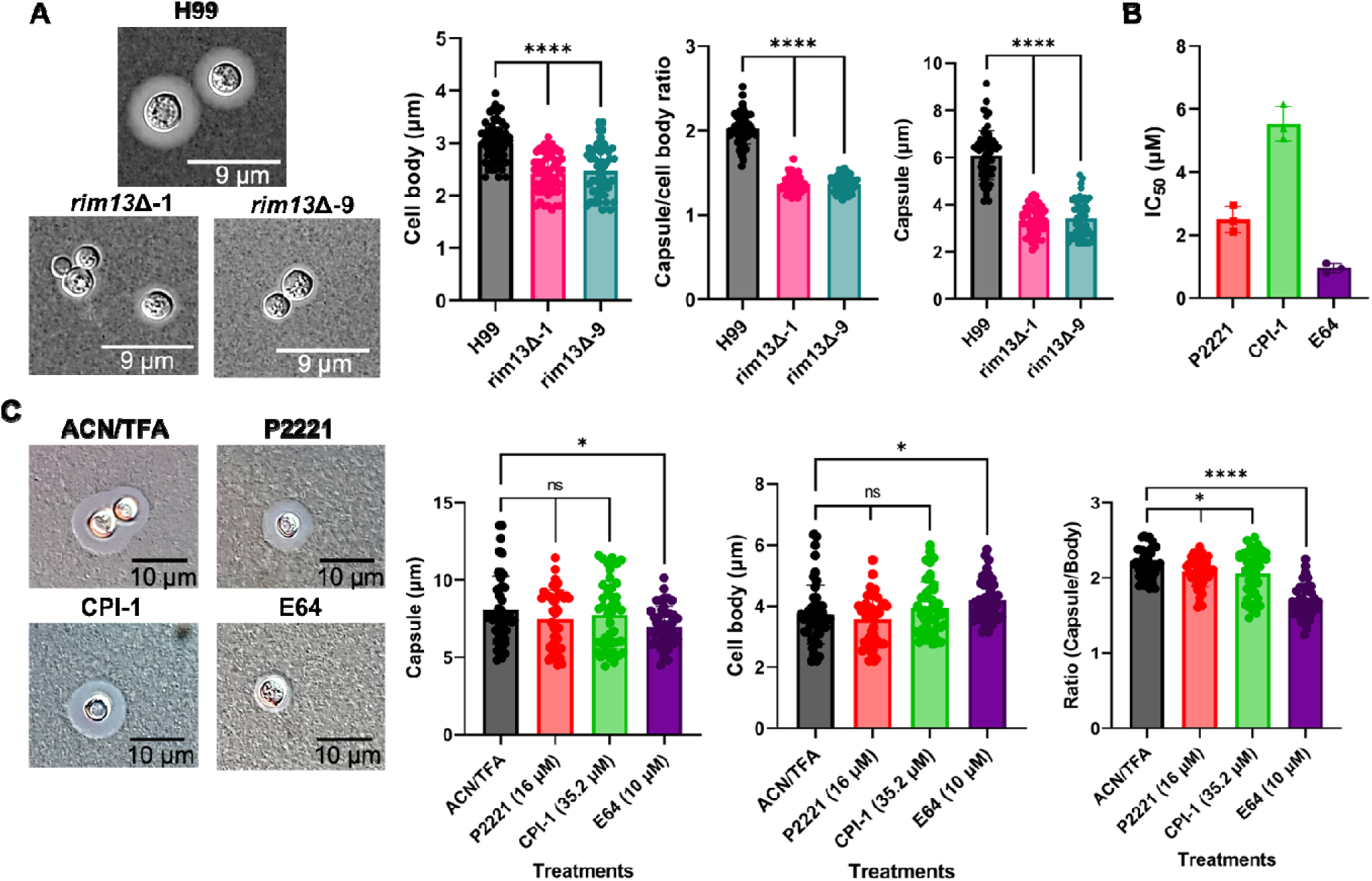
Effect of Rim13 disruption and inhibition on capsule production and fungal cell body size. **A)** Effect of *rim13* deletion on capsule production. **B)** Inhibitory effects of P2221, CPI-1, and E64 on Rim13 enzymatic activity. **C)** Effect of P2221, CPI-1, and E64 on capsule production. Solvent (final 1% ACN/ 0.003% TFA) was used as a control. Experiments were performed using at least three biological replicates. Error bars indicate standard deviation. Statistical analysis was performed using a Welch’s t-test against the solvent treated sample. *: p < 0.05, **: p < 0.01, ***: p < 0.001, ****: p < 0.0001. Statistical figures were prepared using GraphPad Prism 9 (https://www.graphpad.com/). Microscopy images were obtained using a DIC microscope with a 63X oil-based lens.

Given the role of Rim13 in capsule production, we then assessed the ability of P2221 and CPI-1to reduce capsule formation. Interestingly, although capsule and cell size reduction were not significant (p > 0.05), the ratio of capsule/cell body size was significantly decreased upon treatment with P2221 and CPI-1 at 16 and 35 µM, respectively (Fig. 3C). As a control, we observed that 10 µM of E64 significantly reduced the capsule size (p < 0.05) and the ratio to capsule/cell body size; however, an increase in cell body size was observed. Although the reduction in capsule is not as dramatic for fungal cells treated with P2221 and CPI-1 compared to E64, the potential of these computationally derived inhibitors to target the cryptococcal peptidase and disrupt a key virulence factor is evident.

### May1 inhibitors enhance alveolar macrophage clearance

To evaluate the efficacy of our computationally designed peptidase inhibitors through modulation of May1 activity, we focused on the role of secretion of the target peptidase in alveolar macrophage survival. May1 is an aspartic peptidase that allows cryptococcal cells to survive in highly acidic environments, such as the phagolysosome ^56^. First, we aimed to assess the role of *may1* (CNAG_05872) on *C. neoformans* survival within alveolar macrophages by generating a deletion in the H99 background (Fig. S6B, S7B). We observed a significant reduction in fungal cell survival associated with enhanced pathogen clearance by quantifying changes in colony forming unit (CFU) counts for the *may1*Δ strains compared to WT (Fig. 4A). Additionally, we confirmed the role of May1 in growth of *C. neoformans* within low pH environments using *in vitro* plate assays at pH 3.5 and 5.5. As expected, cryptococcal *may1* deletion strains showed a considerable growth defect at pH 3.5 compared to WT but not at pH 5.5 (Fig. 4B). Moreover, we expressed, purified, and confirmed production of FLAG-tagged May1 (Fig. S8B) and assessed *in vitro* inhibitory activity of the two prioritized putative inhibitors, API-1 and API-2. We observed that API-1 and API-2 reduced activity of May1 by 50% at concentrations of 53 and 20 µM, respectively (Fig. 4C). Notably, pepstatin-A (PEP-A), a general aspartic inhibitor and assay control, showed a comparable potency to API-2 (IC_50_ ∼ 25 µM). Further, we evaluated the effect of these two inhibitors on the ability of macrophages to clear phagocytized cryptococcal cells. Both, API-1 and API-2 significantly enhanced fungal clearance by macrophages at 10.6 µM and 12 µM, respectively (Fig. 4D). Interestingly, PEP-A did not cause a significant reduction of fungal burden when used under comparable concentrations and conditions. Together, these data support the ability of API-1 and API-2 to inhibit May1 and impair the ability of *C. neoformans* survival within macrophage using an *in vitro* model of infection.

**Figure 4:**
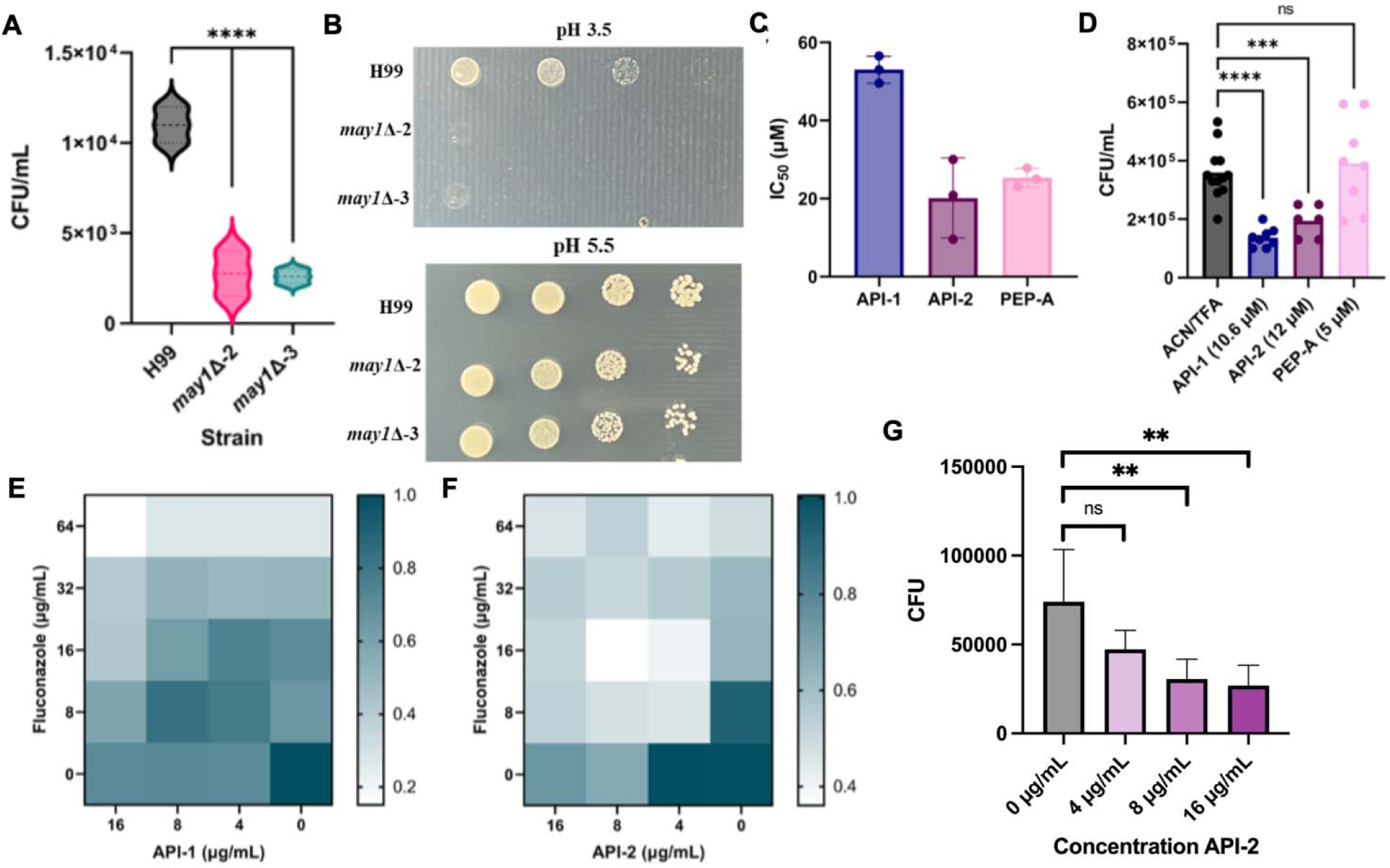
Effect of May1 disruption and inhibition on BALB/c macrophage *C. neoformans* clearance *in vitro* and fluconazole susceptibility. **A)** Effect of *may1* deletion on fungal survival within macrophage as quantified by CFU counts. **B)** Effect of *may1* deletion on *C. neoformans* survival at pH 3.5 and 5.5. **C)** Inhibitory effect of API-1 and API-2 on May1 enzymatic activity. **D)** Effect of API-1, API-2 and PEP-A on macrophage *C. neoformans* clearance. **E** and **F**) Synergy analysi of checkerboard assay of API-1 and API-2 with fluconazole, respectively. G) CFU counts of LE *C. neoformans* cells following co-culture with macrophage and treatment with API-2 at concentrations from 0 to 16 µg/mL and fluconazole at 64 µg/mL. Scale bar measured OD_600nm_. Scale bar measures OD_600nm_. Experiments were performed using a minimum of three biological replicates. Error bars indicate standard deviation. Statistical analysis was performed using a One-Way ANOVA with Dunnett’s multiple comparisons test. *: p < 0.05, **: p < 0.01, ***: p < 0.001, ****: p < 0.0001. Figures were prepared using GraphPad Prism 9 (https://www.graphpad.com/).

### May1 inhibitors influence resensitization of a fluconazole resistant strain to the antifungal

To build on the promise and impact of aspartic peptidases on fungal virulence and a known connection between secreted aspartic peptidases and fluconazole resistance ^57^ (the mainstay treatment option against invasive cryptococcal infections with rising rates of resistance ^58^), we evaluated the efficacy of API-1 and API-2 toward fluconazole re-sensitization. Here, we tested the effect of both candidates on a fluconazole-resistance strain of *C. neoformans* under acidic conditions (YNB pH 3-5) ^35,59^. As observed in our previous experiments, the lab-evolved (LE) strain showed a high degree of resistance towards fluconazole, characterized by a MIC_50_ of 64 µg/mL ^35^. API-1 (at 16 µg/mL) decreased the MIC_50_ of fluconazole 4-fold from 64 to 16 µg/mL and API-2 (at 16 µg/mL) decreased the MIC_50_ of fluconazole 8-fold from 64 to 8 µg/mL (Fig. 4E, 4F). These reductions represent synergistic interactions between API-1 and API-2 at 4 µg/mL toward fluconazole at 8 µg/mL with FICI values of 0.375 and 0.187, respectively. We also tested the effect of API-2 to reduce fungal burden of LE strain in macrophages following treatment with fluconazole and the inhibitor. We observed a significant reduction in survival of the fluconazole resistant cryptococcal strain in the presence of 8 and 16 µg/mL API-2 and 64 µg/mL fluconazole (Fig. 4G). Overall, these data demonstrate resensitization of fluconazole resistant strains to the antifungal, prompting enhanced fungal cell clearance from macrophage and recovery of drug efficacy.

### CnMpr1 inhibitors reduce BBB crossing of C. neoformans

To evaluate the efficacy of our computationally designed peptidase inhibitors through modulation of CnMpr1 activity, we focused on the role of the target peptidase in BBB crossing. Given that CnMpr1 is a secreted metallopeptidase used by *C. neoformans* for brain invasion ^60^, we assessed the capacity of our predicted metallopeptidase inhibitors against BBB crossing using an *in vitro* model. First, we generated a gene deletion strain for *cnmpr1* (CNAG_04735) in the *C. neoformans* H99 background and confirmed its role in BBB crossing using an *in vitro* endothelial cell model. We observed a significant reduction in BBB crossing in the mutant strain compared to WT (Fig. 5A). Next, we attempted to express and purify FLAG-tagged CnMpr1 for *in vitro* assays; however, we were not successful and alternatively, we used a commercially available M4 metallopeptidase structurally related to CnMpr1 (thermolysin) a as a model for the enzymatic assays ^19^. Notably, thermolysin (UniProt: P00800) shares 17% sequence identity with CnMpr1 with a Z-score of 23.7 (Fig. S9) ^61^ with interaction predictions compared for each peptidase and the designed inhibitors showing higher predicted binding affinity (ipTM and LIA scores) for the inhibitors toward CnMpr1 (Fig. S10). Treatment of thermolysin with the predicted inhibitors showed low inhibitory activity for MPI-1 and MPI-2 with IC_50_ values of 118 and 90 µM, respectively, compared to stronger inhibitory activity for MPI-3 and MPI-4 with IC_50_ values of 9.8 and 1 µM, respectively (Fig. 5B). Similarly, ethylenediaminetetraacetic acid (EDTA), a known non-specific divalent metal chelator and metallopeptidase inhibitor, showed an IC_50_ of approx. 0.1 µM against Thermolysin ^62^.

**Figure 5:**
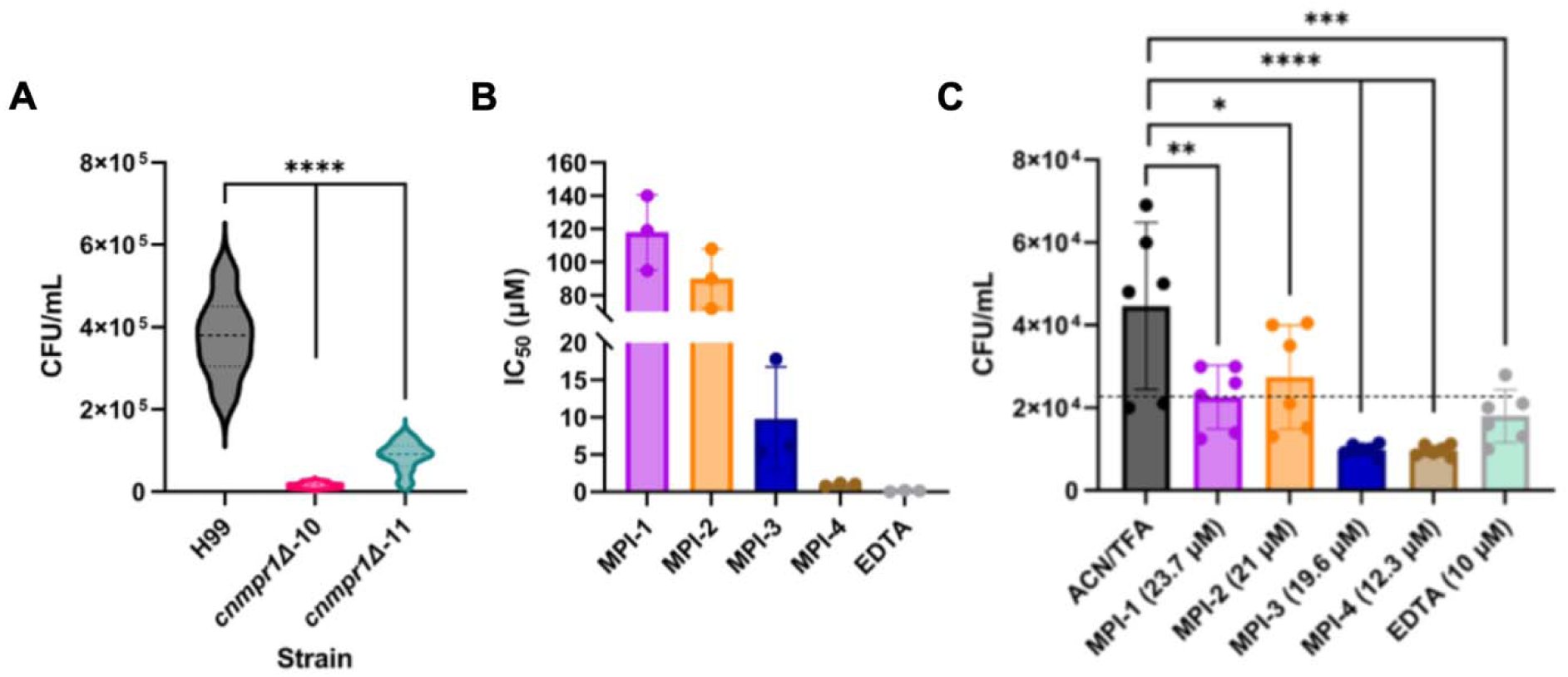
Effect of *cnmpr1* disruption or chemical inhibition on *C. neoformans* BBB crossing. **A)** Effect of gene deletion of *cnmpr1* in BBB crossing *in vitro* assay. **B)** Inhibitory effect of candidates and EDTA on Thermolysin enzymatic activity. **C)** Effect of MPI-1, −2, −3, and-4 and EDTA on *C. neoformans* BBB crossing using an *in vitro* model. Dash line indicates 50% inhibition compared to the control. Experiments were performed using at least three biological replicates. Error bars indicate standard deviation. Statistical analysis was performed using a One-Way ANOVA with a Dunnett’s multiple comparisons test. *: p < 0.05, **: p < 0.01, ***: p < 0.001, ****: p < 0.0001. Statistical figures were prepared using GraphPad Prism 9 (https://www.graphpad.com/).

Next, we aimed to assess if the inhibitors impacted *C. neoformans* BBB crossing *in vitro* upon treatment. All inhibitors showed significant (p < 0.05) reduction of *C. neoformans* H99 BBB crossing compared to the control (solvent) based on CFU counts (Fig. 5C). In this context, while MPI-1 and MPI-2 inhibited approximately 50% of the BBB crossing, MPI-3 and MPI-4 achieved more than 75% inhibition relative to the control. Likewise, we also observed that EDTA significantly inhibited BBB crossing. Together, these data support the roles of MPI-3 and MPI-4 to inhibit the metalloprotease enzyme and impair the ability of *C. neoformans* to cross the BBB within an *in vitro* model.

### Peptidase inhibitors reduce fungal virulence without causing host cytotoxicity

Given the impact of the inhibitors toward capsule/cell size ratio, macrophage clearance, fluconazole resistance, and BBB crossing, we next aimed to assess the ability of the inhibitors to reduce virulence of *C. neoformans*. Since the capsule is needed to survive inside the phagolysosome and CnMpr1 is also involved in macrophage evasion, we tested if all inhibitors could reduce fungal survival and enhance macrophage clearance^12^. To mimic the natural course of the infection, we allowed the cryptococcal cells to be phagocytosed before the application of inhibitors at a concentration of 10 µg/mL. We observed that, except for MPI-1 and MPI-2, the remaining inhibitors significantly reduced fungal survival within macrophage (Fig. 6A). We also assessed the cytotoxic effect of the inhibitors against mammalian cells by quantifying the release of lactate dehydrogenase by macrophages upon inhibitor treatment. We observed no significant difference in LDH release upon inhibitor treatment compared to the untreated controls (Fig. 6B).

**Figure 6:**
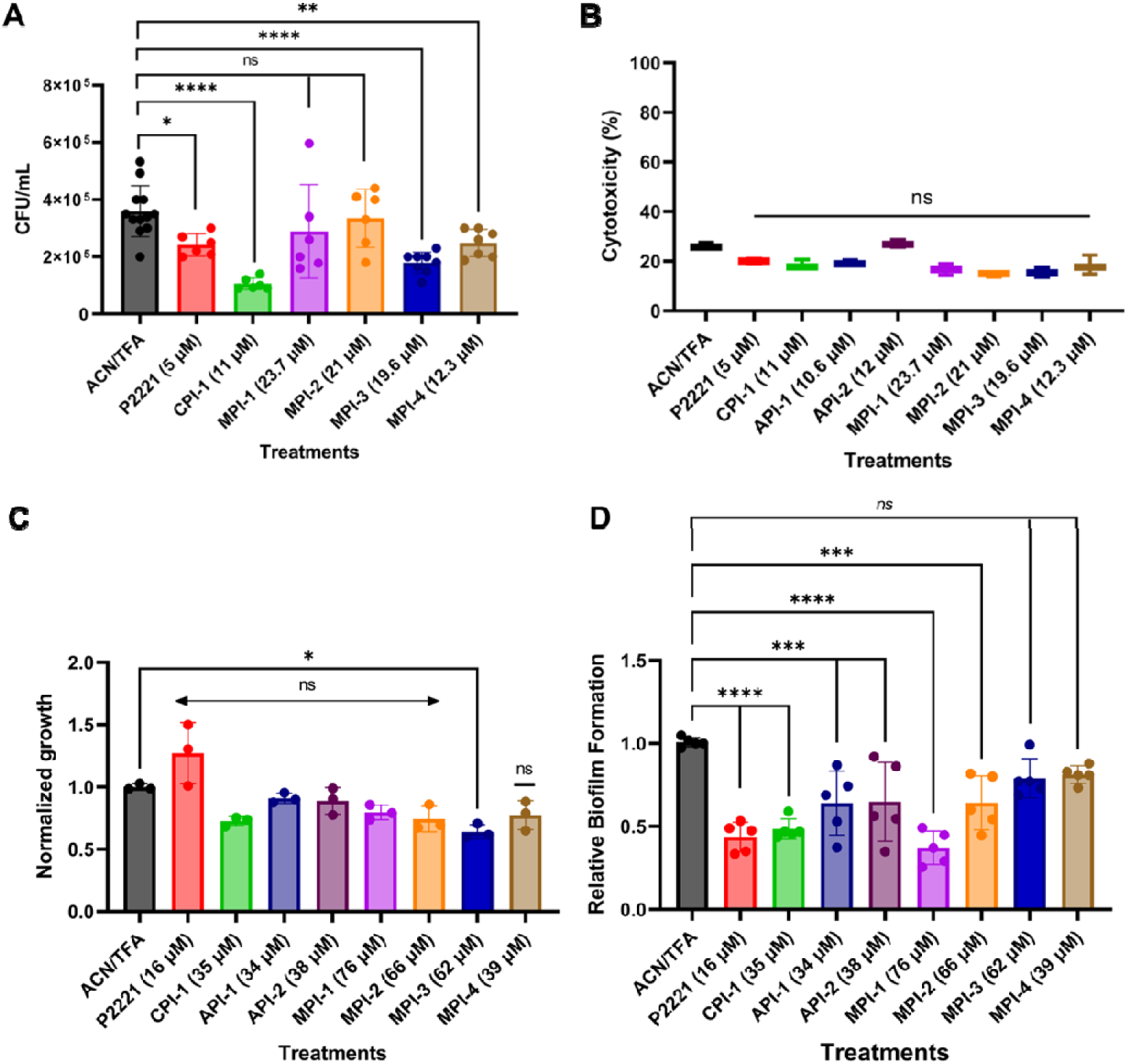
Effect of peptidase inhibitors on *C. neoformans* pathogenicity using *in vitro* models. **A)** BALB/c macrophage clearance of *C. neoformans in vitro* at 10 µg/mL. **B)** Cytotoxic effect of inhibitors against BALB/c macrophages at 10 µg/mL. **C)** Effect on fungal growth in YNB media at 32 µg/mL. **D)** Biofilm formation assay. Growth and biofilm formation were normalized to the control (ACN/TFA). All candidates were tested at a final concentration of 32 µg/mL and solvent, (final 1% ACN/ 0.003% TFA) was used as control. Experiments were performed using at least three biological replicates. Error bars indicate standard deviation. Statistical analysis was performed using a One-Way ANOVA with a Dunnett’s multiple comparisons test. *: p < 0.05, **: p < 0.01, ***: p < 0.001, ****: p < 0.0001. Figures were prepared using GraphPad Prism 9 (https://www.graphpad.com/).

Next, we evaluated the potential fungistatic (inhibition of growth) or fungicidal (killing of fungal cells) effects of each inhibitor against *C. neoformans* H99 growth *in vitro*. Here, we applied 32 µg/mL for each inhibitor, which is higher than the MIC_50_ breakpoints for most antifungal agents used to treat fungal infections ^63^. We did not observe a significant change in fungal growth upon inhibitor treatment except for the metallopeptidase inhibitor, MPI-3, which significantly reduced fungal growth by approx. 25% (Fig. 6C). In addition, none of the inhibitors appeared to have fungicidal effects toward *C. neoformans*, indicating that inhibitor treatment is influencing virulence factor production instead of fungal growth, leading to lower selective pressure toward resistance ^9,13,64^. Next, given the clinical importance of biofilms and their connection to capsule production and brain invasion ^65,66^, we investigated anti-biofilm capabilities of the peptide inhibitors. We observed that, all candidates except MPI-3 and MPI-4, significantly reduced the formation of cryptococcal biofilms when applied at 32 µg/mL (Fig. 6D). These results demonstrate the clinical potential of the peptidase inhibitors in reducing fungal burdens and biofilms while limiting the host cytotoxicity and the evolution and antifungal resistance.

### CPI-1, API-1 and MPI-3 enhance fluconazole activity in vivo

Based on the antivirulence and inhibitory properties of the computationally predicted inhibitors, along with their potential clinical value, we aimed to evaluate inhibitor effectiveness *in vivo*. Here, we tested the ability of one inhibitor from each enzymatic family (i.e., CPI-1, API-1 and MPI-3) to reduce the virulence of *C. neoformans* in a *Galleria mellonella* (i.e., greater wax moth) larvae model ^48^. Although this model lacks the adaptive immune system of mammalian models (e.g., mice), it allows researchers to quickly assess the potential of new compounds before moving into more complex and expensive *in vivo* assays. Notably, these assays were performed over 5 days post inoculation (dpi) with full assessment of inhibitor efficacy performed at 3 dpi, when larvae were healthy and consistent differences in survival were observed.

First, we evaluated the toxicity of the inhibitors with or without fluconazole and found no significant difference in inhibitor toxicity compared to the solvent (PBS) (Fig. 7A). Then, to imitate a clinical infection, we infected the larvae with fungal cells before applying treatment and we observed that while fluconazole (positive control) reduced the infection significantly, none of the inhibitors alone significantly reduced fungal virulence (Fig. 7B). Finally, given our observation of additive effects between inhibitor and fluconazole in a macrophage model, we tested if the inhibitors could enhance fluconazole antifungal activity *in vivo*. We observed a significant difference in larvae survival upon combinatory treatment with inhibitor and fluconazole at 3 dpi compared to fluconazole alone or the untreated controls (Fig. 7C). By 4 and 5 dpi, these differences were reduced, suggesting that inhibitor stability or declining larval health may have confounding effects on inhibitor efficacy. Overall, these results demonstrate the potential of the peptidase inhibitors as enhancers of approved antifungal agents in combinatory therapies.

**Figure 7:**
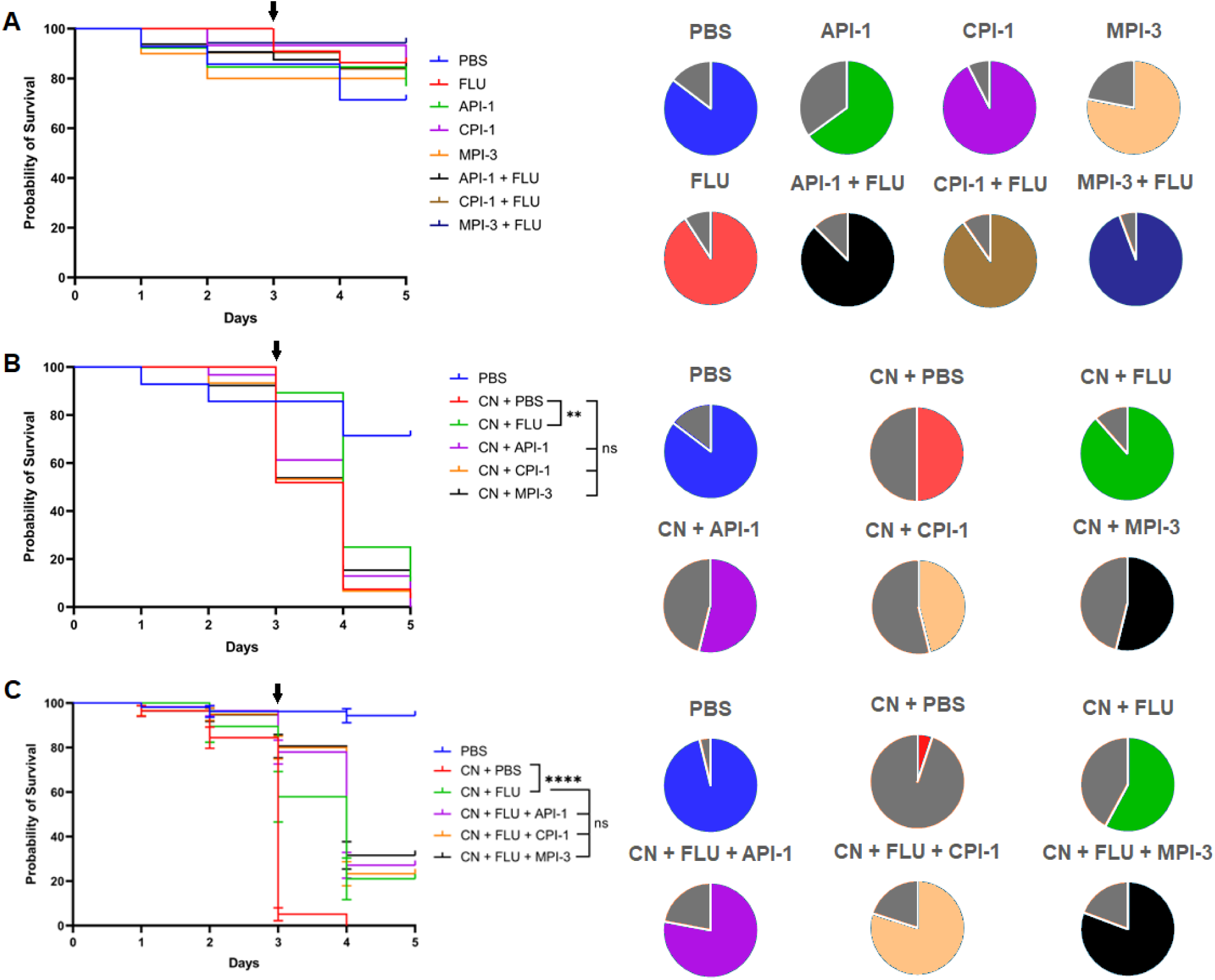
Effect of inhibitors on survivability of *G. mellonella* larvae after infection with *C. neoformans* H99. **A)** Host toxicity of CPI-1, API-1 and MPI-3 with or without fluconazole at 10 mg/kg in absence of fungal infection. **B)** Effect of CPI-1, API-1 and MPI-3 upon infection with *C. neoformans* and treatment with the inhibitors. **C)** Effect of CPI-1, API-1 and MPI-3 upon infection with *C. neoformans* with fluconazole at 10 mg/kg. Between 40 and 60 larvae were used per condition. CN stands for *C. neoformans* and FLU for fluconazole. Data for pie charts were obtained at 3 dpi. Color in pie charts indicate percentage of alive larvae at 3 dpi while grey indicates dead larvae. Error bars indicate standard deviation. Statistical analysis was performed using a log-rank (Mantel-Cox) test. *: p < 0.05, **: p < 0.01, ***: p < 0.001, ****: p < 0.0001. Figures were prepared using GraphPad Prism 9 (https://www.graphpad.com/).

### CPI-1 inhibitor reduces cryptococcal polysaccharide capsule through altered cell wall synthesis and integrity

To define the downstream effects of the inhibitors and elucidate their anti-virulence mechanisms, we performed mass spectrometry-based proteomics on the treated and untreated samples. As Rim13 is an intracellular target and we aimed to assess the impact of a Rim13 inhibitor toward the target and downstream effects, we treated *C. neoformans* H99 cells with CPI-1 under capsule-inducing conditions (LIM and 37 °C). Here, we observed a significant shift in the fungal proteome upon treatment, with identification of 4201 proteins (52% proteome [UP000010091] coverage) with 244 proteins showing significantly higher abundance in the untreated samples vs. 243 proteins with significantly lower abundance upon CPI-1 treatment (Fig. 8A). Similarly, a principal component analysis (PCA) revealed a clear separation of samples after treatment with CPI-1 (Fig. 8B). Moreover, to shed light on the effects of Rim13/RIM pathway inhibition via CPI-1 treatment, we focused our analysis on proteins with reduced abundance following treatment. Based on Gene Ontology analysis, lower abundance proteins were grouped into two main biological processes: i) translation (e.g., ribosome biogenesis, RNA processing and gene expression) and ii) metabolic processes (e.g., members of the electron transfer chain) (Fig. 8C; Fig. S11). Notably, we did not identify Rim13 within our analysis, suggesting that the abundance of this protein was below the limit of detection under the tested conditions.

**Figure 8:**
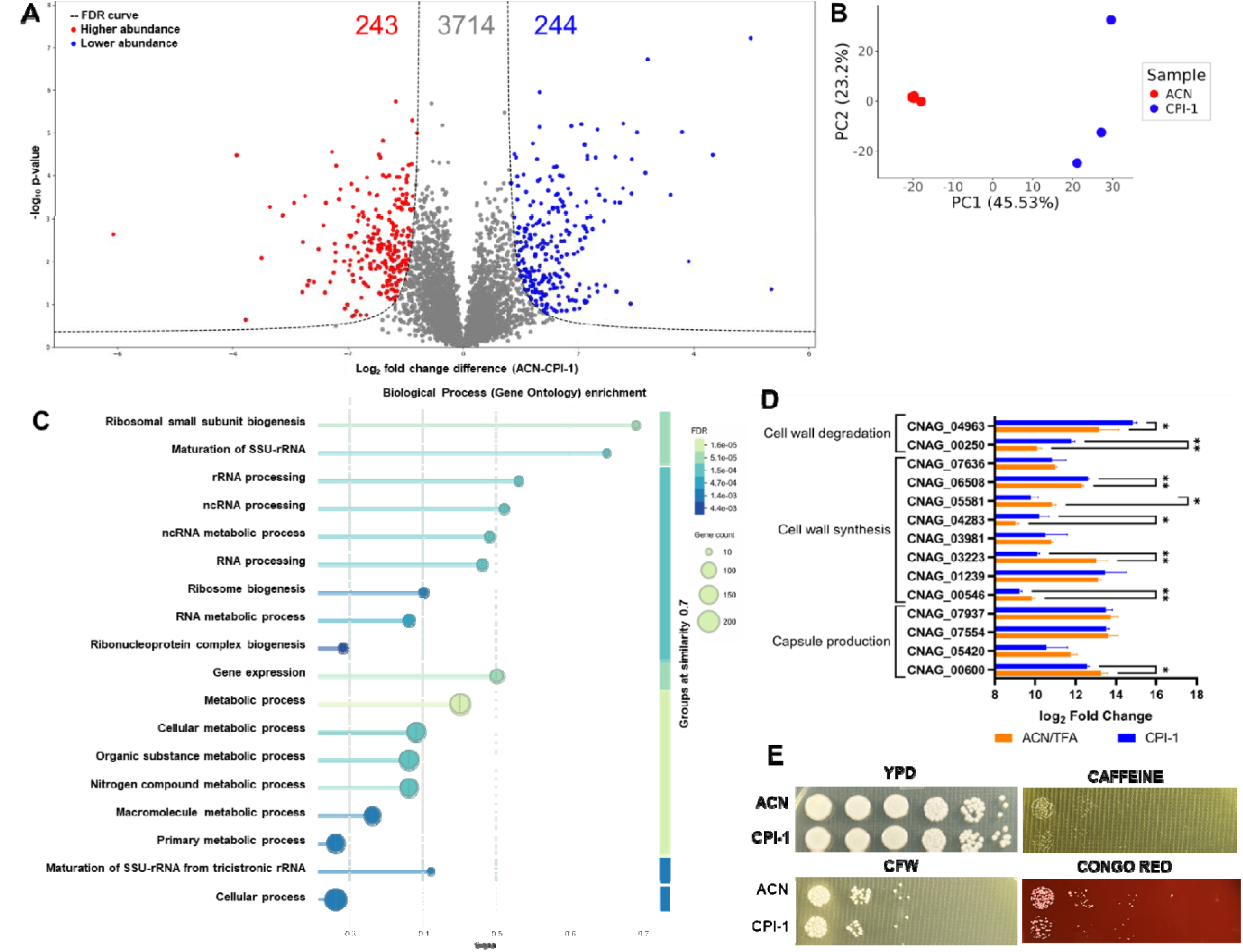
Proteomic analysis of CPI-1 effects on *C. neoformans* proteome. **A)** Volcano Plot showing significant changes in protein abundance. Values on the left side (red dots) have higher abundance in untreated samples and on the right (blue dots) higher abundance upon treatment with CPI-1. Statistical analysis = Student’s t-test, p-value < 0.05; FDR = 0.05; S_0_ = 1. Figure was prepared using Python. **B**) Principal Component Analysis (PCA) showing two main sources of variation. Samples treated with ACN (control) as red circles and with CPI-1 in blue. **C)** Biological Process Gene Ontology enrichment of 244 proteins with significantly lower abundance upon treatment. Analysis was performed using the STRING DB and *C. neoformans* H99 proteome (UP000010091) as background. Biological processes are sorted by signal (harmonic mean between the observed/expected ratio and - log[FDR]) and grouped with a similarity of 0.7. **D)** Comparison of protein abundance of 14 genes involved in cell wall degradation, cell wall synthesis and capsule production. Gene numbers are sorted from high to low within each category. **E)** Effect of CPI-1 on *C. neoformans* H99 cells under cell wall stress. CFW = calcofluor white. Error bars indicate standard deviation. Statistical analysis was performed using a Student T-test comparing abundances within each gene. For visual simplicity, only comparisons with a p-value lower than 0.05 are shown. *: p < 0.05, **: p < 0.01. Figures were prepared using GraphPad Prism 9 (https://www.graphpad.com/).

Since we predict that CPI-1 targets Rim13, which activates the transcriptional factor Rim101 involved in cell wall and capsule formation, we specifically compared changes in protein abundance of 14 genes important for these processes (Fig. 8D; Table S10). We observed that two proteins (CNAG_00250 and CNAG_04963) involved in cell wall degradation were significantly higher in abundance following treatment with CPI-1 compared to the control, supporting compensatory processes that may disrupt capsule production through changes in the cell wall. On the other hand, three (CNAG_00546, CNAG_05581, and CNAG_03223) out of eight proteins involved in cell wall synthesis showed reduced abundance upon treatment, and two proteins (CNAG_04283 and CNAG_06505) showed increased abundance, highlighting the complexity of this pathway. Importantly, CNAG_00600 (encodes for Cap60), which is critical for sporulation, capsule formation and virulence showed a significant reduction in CPI-1 treated cells ^67,68^. Lastly, to validate our proteomics results, we assessed the effect of CPI-1 on cryptococcal cells in the presence or absence of cell wall stressors *in vitro*. Notably, we observed slight reductions in cell wall integrity and stability upon CPI-1 treatment in the presence of caffeine (cell wall integrity stressor via TOR/MAPK pathway^69,70^), Congo Red (binds β-glucans in fungal cell walls^71^), and calcofluor white (binds to chitin^72^) compared to no stress (YPD) conditions (Fig. 8E). Together, proteomics profiling integrated with phenotypic assays demonstrate remodeling of the *C. neoformans* cell wall upon treatment with CPI-1 and supports its predicted mode of action through disruption of Rim13 and its related regulatory roles driving capsule synthesis and cell wall maintenance and stability.

## Discussion

Fungal infections, such as those caused by *C. neoformans*, are increasing in incidence and mortality, which are accompanied by rising rates of antifungal resistance^3,4^. Virulence factors, important for infection but not growth, could be an answer to drug development challenges. For instance, by targeting virulence traits, it is possible to lower the pathogens’ defenses and enhance natural immune responses while limiting the selective pressure towards resistance. Here, we developed a computational pipeline coupled with structure-based design to prioritize putative inhibitors of three virulence-related peptidases in *C. neoformans:* Rim 13, May1, and CnMpr1. The best performing thresholds were applied to screen inhibitor candidates from natural sources against *C. neoformans* peptidase targets, followed by inhibitory assays, phenotypic assessments and alignment with genetic deletion strains, along with *in vitro* and *in vivo* virulence evaluation and mechanism of action studies. Overall, the designed peptidase inhibitors exhibited potent antifungal activity without harming mammalian cells, establishing a predictive framework for rational scaffold design of next-generation antifungals that disarm the pathogen to enable immune-mediated clearance.

We observed statistically significant differences between TP and TN complexes across all scoring metrics (ipTM, LIS, and LIA), with mean TPRs at optimal thresholds ranging from 0.891–0.934 and corresponding FPRs of 0.043–0.087, indicating that nearly all true peptidase–inhibitor pairs can be identified with <10% false positives. Classification accuracy was higher for the peptidase–protein dataset than for peptidase–peptide datasets, likely reflecting the predominance of protein–protein complexes in the AFM training data. Across evaluation datasets, ipTM and LIS showed the strongest performance; however, LIS consistently achieved a slightly lower FPR at comparable TPRs for peptide–peptidase complexes, while both metrics performed equivalently for protein datasets, supporting LIS as a more reliable indicator for peptide–peptidase prediction. This difference likely arises from metric design: ipTM assesses confidence in global chain positioning, whereas LIS and LIA are derived from predicted alignment error (PAE) at the interaction interface ^25,27^. LIA reflects the interface area below a predefined PAE threshold and scales with interface size, while LIS incorporates the average inverted PAE within this region, increasing only with lower PAE values that better reflect higher-confidence, affinity-relevant interactions. Further, we developed the pipeline with AFM as it provides comparable protein-protein complex predictions to AlphaFold3 with current modeling and has established scoring metric thresholds for protein-inhibitor complex predictions^28^.

We confirmed through gene deletion that disruption of *rim13* markedly impairs capsule production and prioritized the design of two Rim13 inhibitors, P2221 and CPI-1. Both compounds exploit the calpain-like catalytic mechanism of Rim13, which involves formation of an acyl-enzyme intermediate, enabling inhibition through covalent cysteine-targeting warheads ^73^. Structural modeling predicted that P2221 binds the active-site cysteine (C185) via a disulfide bond and engages stabilizing hydrophobic interactions, although its extended structure likely limits intracellular accessibility. To improve pharmacological properties, CPI-1 was designed as a shorter derivative with enhanced binding metrics and improved predicted membrane penetrability. Both inhibitors moderately inhibited recombinant Rim13 *in vitro*; however, IC values may be influenced by the use of a general peptidase substrate, highlighting the need for a Rim13-specific assay ^21^. While both compounds similarly reduced capsule formation, CPI-1 more effectively enhanced macrophage-mediated fungal clearance and impaired biofilm formation likely through disrupted capsule attachment ^74^, and exhibited additive efficacy with fluconazole in the *G. mellonella* model, potentially by increasing drug penetration. Importantly, CPI-1 attenuates fungal virulence rather than viability and displays low host cytotoxicity, suggesting reduced selective pressure for resistance development ^75^. Complementary mass spectrometry-based proteomics revealed broad intracellular effects of CPI-1, with 11% of the detected proteome significantly altered with marked reductions in proteins linked to translation, metabolism, and capsule biosynthesis, consistent with impaired Rim101 activation ^30^. Compensatory changes in cell wall biogenesis proteins likely reflect engagement of alternative regulatory pathways, such as protein kinase A signaling ^14,29,76,77^, while the downregulation of cell wall degradation enzymes further supports disruption of RIM pathway-mediated remodeling. Collectively, these findings validate Rim13 as a druggable virulence regulator and position CPI-1 as a promising lead for antivirulence antifungal development.

A critical barrier to *C. neoformans* infection is the alveolar macrophage, which engulfs pathogens into acidic phagolysosomes for elimination. To survive this hostile environment, *C. neoformans* secretes the pepsin-like aspartic peptidase May1, which degrades host proteins to provide nutrients and dismantle immune defenses, thereby enhancing virulence ^11^. Given its central role in immune evasion, May1 represents an attractive therapeutic target. Using our computational pipeline, we identified two inhibitors of May1, trypsin 3075 and trypsin 735, and leveraged structural comparisons with the May1-pepstatin A complex (PDB: 6R6A) to guide inhibitor optimization. Specifically, substitution of key residues with statine (critical for pepstatin A inhibition ^78^) yielded two optimized peptidomimetics, API-1 and API-2, while preserving flanking residues to maintain active-site engagement. Structural modeling predicted stabilizing interactions within the S1′ and S3 pockets for both inhibitors, with additional hydrophobic interactions unique to API-2 likely accounting for its higher affinity toward May1.

Both API-1 and API-2 inhibited May1 with micromolar IC values; however, these measurements likely underestimate potency due to the use of an oversaturating general substrate, which can mask competitive inhibition effects ^21^. Notably, pepstatin A (despite nanomolar affinity for pepsin-like peptidases ^79^) exhibited comparable inhibition under these assay conditions, supporting the functional relevance of the designed inhibitors. Importantly, both compounds reduced fungal burden within alveolar macrophages, and API-2 further enhanced fluconazole efficacy in acidic conditions and intracellular environments, possibly by limiting extracellular proteolysis and impairing fungal growth. Given the growing threat of antifungal resistance ^80^, these findings are particularly significant, as acidic environments and biofilm formation are known contributors to fluconazole resistance in *C. neoformans* ^81–83^. Consistent with this, both API-1 and API-2 significantly reduced biofilm formation, aligning with the established role of secreted aspartic peptidases in surface adherence ^84^. *In vivo* studies in *G. mellonella* demonstrated additive effects with fluconazole, likely through inhibition of May1 activity within hemocytes ^85,86^. Collectively, these data highlight May1 inhibition as a promising antivirulence strategy that enhances host immunity and existing antifungal therapies while potentially reducing the emergence of resistance.

Although *C. neoformans* can infect immunocompetent hosts, it primarily causes fatal meningoencephalitis in immunocompromised individuals by escaping alveolar macrophages, disseminating systemically, and crossing the BBB ^87^. Preventing BBB traversal represents a critical therapeutic objective. A key mediator of this process is the secreted metallopeptidase CnMpr1, which is essential for BBB crossing and represents an attractive antifungal target. Using two complementary design strategies: incorporation of metal-chelating functional groups and structure-based computational modeling, we developed four CnMpr1 inhibitors (MPI-1–4). Genetic deletion of *cnmpr1* confirmed its requirement for BBB crossing *in vitro*, validating the enzyme as a druggable target. Among the designed compounds, MPI-3 and MPI-4 demonstrated the strongest inhibition of thermolysin (IC ∼1–10 µM), a commercial metallopeptidase surrogate for CnMpr1, likely due to favorable active-site interactions absent in MPI-1 and MPI-2 (IC ∼90–118 µM). Both MPI-3 and MPI-4 contain valine residues predicted to engage a hydrophobic region near the active site and polar groups capable of coordinating the catalytic zinc ion, a hallmark of effective metallopeptidase inhibitors. The slightly reduced potency of MPI-3 relative to MPI-4 may reflect its lack of interactions with the S1′ pocket. While use of a non-homologous commercial enzyme is a limitation, MPI-3 and MPI-4 most effectively reduced BBB crossing, consistent with their superior enzymatic inhibition, though MPI-1 and MPI-2 also significantly impaired BBB traversal, suggesting that even partial inhibition of CnMpr1 is sufficient to disrupt this process. Under neutral pH conditions that promote CnMpr1 secretion^11^, only MPI-3 and MPI-4 reduced intracellular fungal burden within macrophages, likely by limiting extracellular proteolysis or nutrient acquisition ^88^. None of the inhibitors affected fungal growth or exhibited mammalian cytotoxicity. Interestingly, MPI-1 and MPI-2, but not MPI-3 or MPI-4, reduced biofilm formation, potentially through nonspecific chelation of divalent cations, such as Mg² and Ca² by sulfonamide groups, a mechanism observed with general metal chelators like EDTA ^89^. Notably, MPI-3 exhibited additive effects with fluconazole in the *G. mellonella* model, likely through impaired nutrient acquisition and reduced immune protein degradation ^88,90^. Collectively, MPI-3 and MPI-4 emerge as promising leads for targeting BBB invasion, *C. neoformans*’ most lethal virulence trait, and provide a strong foundation for future therapeutic development.

## Conclusion

*Cryptococcus neoformans* is a human fungal pathogen that infects and kills thousands of immuno-compromised individuals every year. Although treatment is available, due to several factors, antifungal resistance is becoming more prevalent, especially in low-income countries where the need for effective therapy is greatest. By attacking its virulence factors, we aim to disarm the pathogen, enhancing natural defenses or efficacy of other antifungal agents, which lowers the risk of promoting resistance. Here, we developed an AFM-based pipeline for the discovery of novel peptidase inhibitors and, combined with structure-based rational design, we prioritized and evaluated eight inhibitors against three virulence-related peptidases in *C. neoformans*. These new agents not only affect important virulence factors, such as polysaccharide capsule production and biofilm formation, they also reduce BBB crossing, enhance the alveolar macrophage fungal clearance *in vitro,* and increase fluconazole effectiveness *in vivo* without targeting host cells. Together, these promising findings provide an integrative computational-experimental scaffold for antifungal drug development with promising efficacy and clinical relevance.

## Acknowledgements

The authors thank members of the Geddes-McAlister lab for helpful discussions and constructive comments on the study. We also thank the members of Dr. Tariq Akhtar’s lab, including Dr. Kelly Boddington and Dr. Eric Soubeyrand, for technical assistance to evaluate peptide purity.

## Author Contributions

D.G.-G., J.H., R.S.P., A.H.-W. & J.G.-M. conceptualized and designed the study. D.G.-G., J.H., E.O.-A., P.A.V., A.H.-W. & J.G.-M. contributed to computational pipeline development and application. D.G.-G., N.C., O.R., M.W., A.D., and M.A., performed *in vitro* experiments. D.G.-G., M.J.M., A.D., and J.G.-M. designed and/or performed *in vivo* experiments. G.W. synthesized peptides. D.G.-G., M.W., J.D., S.N.S., and J.G.-M. designed and performed mass spectrometry experiments. D.G.-G., J.H., M.W., A.D., M.J.M., J.D., S.N.S., P.A.V., M.A., A.S., A.H.-W., and J.G.-M. analyzed data. D.G.-G., J.H., and J.G.-M. generated figures and tables. D.G.-G., J.H., A.H.-W., and J.G.-M. wrote the first manuscript draft. All authors contributed to manuscript preparation and have read and approved the submitted manuscript.

## Data availability

The proteomics datasets are available through PRIDE Proteomics Exchange.

Project Accession: PXD072145

Token: TJRUaDOzGGzx

## Computational resources

Hardware requirements for the computational analyses were provided by the Graham High-Performance Computing Cluster (Compute Canada). One Nvidia GPU (V100, A100, or P100) was allocated for each AFM and AF2 prediction. AlphaPulldown was executed using a Docker container image (gallardoalba/alphapulldown:0.30.7) to generate structures of peptidase-inhibitor complexes and AlphaFold 2.3 was used to generate the solo peptidase structures. Alpha-Pulldown and AlphaFold were run using five recycling steps with default parameters unless otherwise specified. Job submission and resource allocation were managed using SLURM workload manager. Scripts used to run the pipeline can be found at https://github.com/HamSlice7/Peptidase-Inhibitor-Screening-Pipeline/tree/main.

