## Supplementary material for "A structure-guided pipeline yields peptide inhibitors that disarm fungal peptidase-driven virulence and resistance": Supp fig

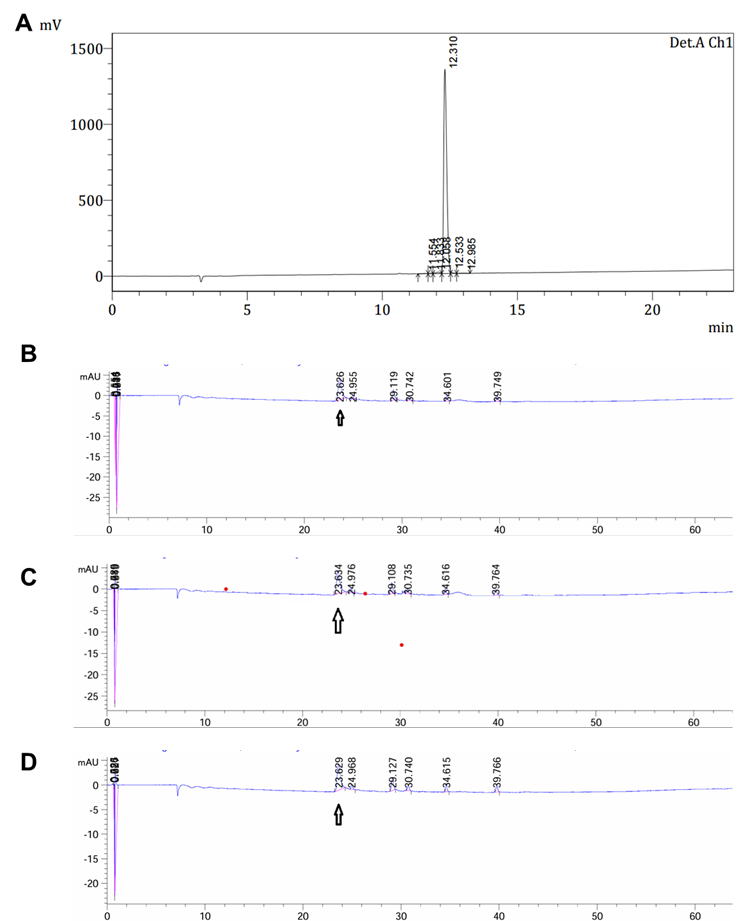

**Figure S1:** **HPLC analysis of peptide synthesis.** **A:** P2221. **B:** CPI-1. **C:** API-1. **D**: API-2. Purity was assessed with a SHIMADZU Inertsil ODS-SP(4.6*250mm*5um) column across a 60 min gradient of Buffer A (0.1% TFA in 100% water) to Buffer B (0.1% TFA in 100% acetonitrile) and OD_220nm_ measured.

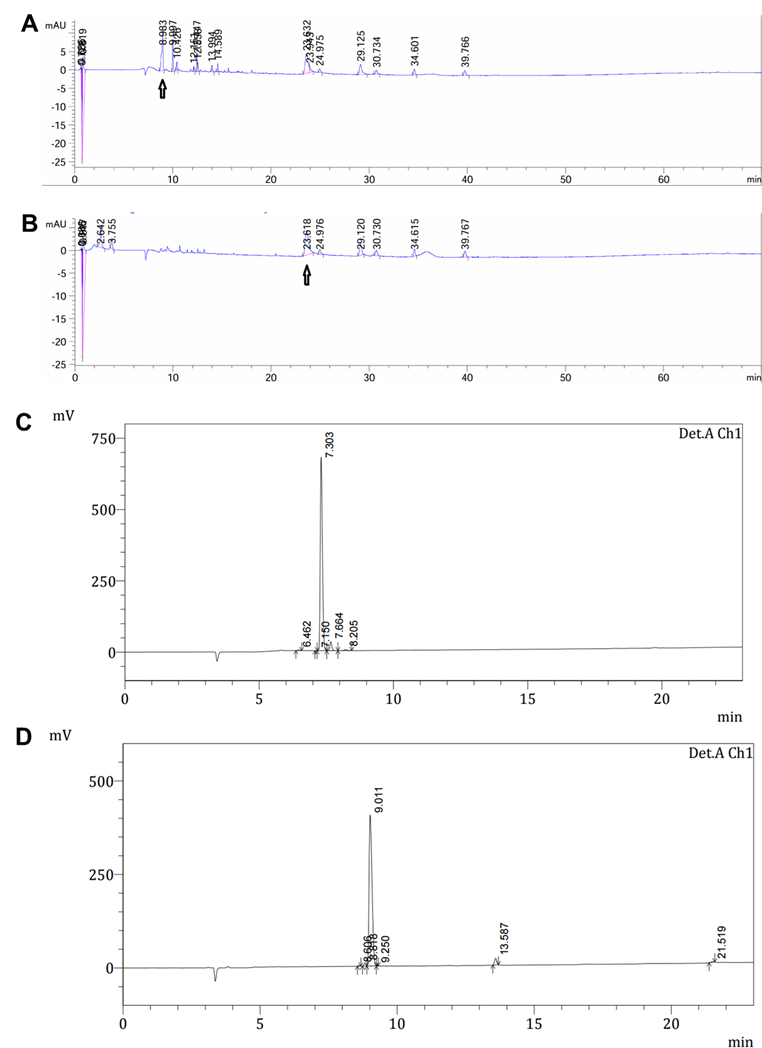

**Figure S2: HPLC analysis of peptide synthesis. A:** MPI-1. **B:** MPI-2. **C:** MPI-3. **D:** MPI-4. Purity was assessed with a SHIMADZU Inertsil ODS-SP(4.6*250mm*5um) column across a 60 min gradient of Buffer A (0.1% TFA in 100% water) to Buffer B (0.1% TFA in 100% acetonitrile) and OD_220nm_ measured.

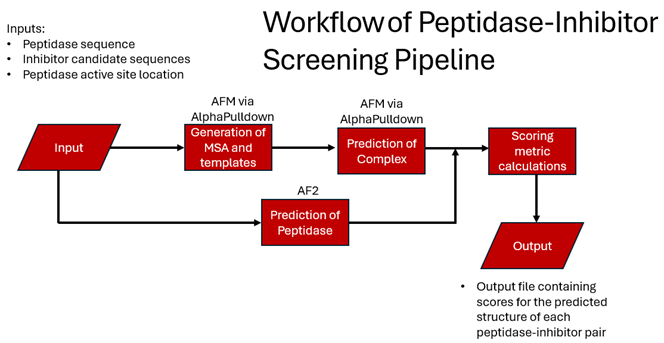

**Figure S3:** **Computational pipeline development.** Computational workflow to screen peptidase-inhibitor complexes. AFM = AlphaFold Multimer; MSA = Multiple Sequence Alignment.

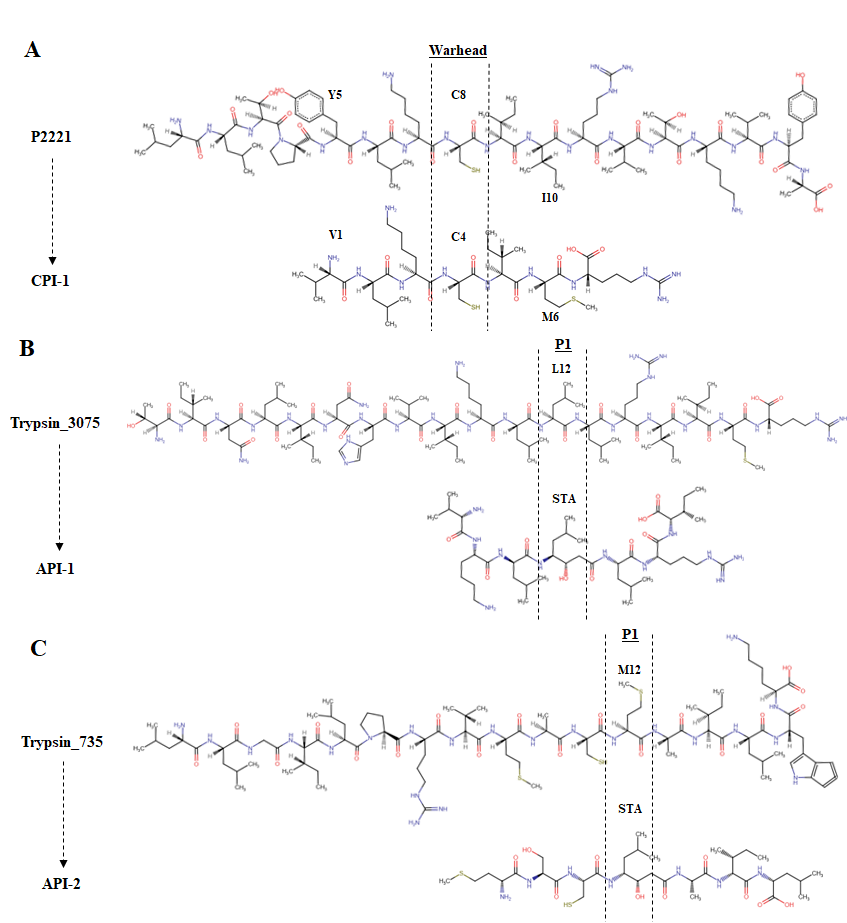

**Figure S4:** **Structure based design of inhibitors against Rim13 and May1.** **A**: structure of P2221 and CPI-1, Rim13 inhibitors. **B**: design of API-1 based on Trypsin_3075 as May1 inhibitor. **C**: design of API-2 based on Trypsin_735 as May1 inhibitor. Labeled amino acids where mutated during the optimization stage. STA stands for statine. P1 position was highlighted with dashed lines.

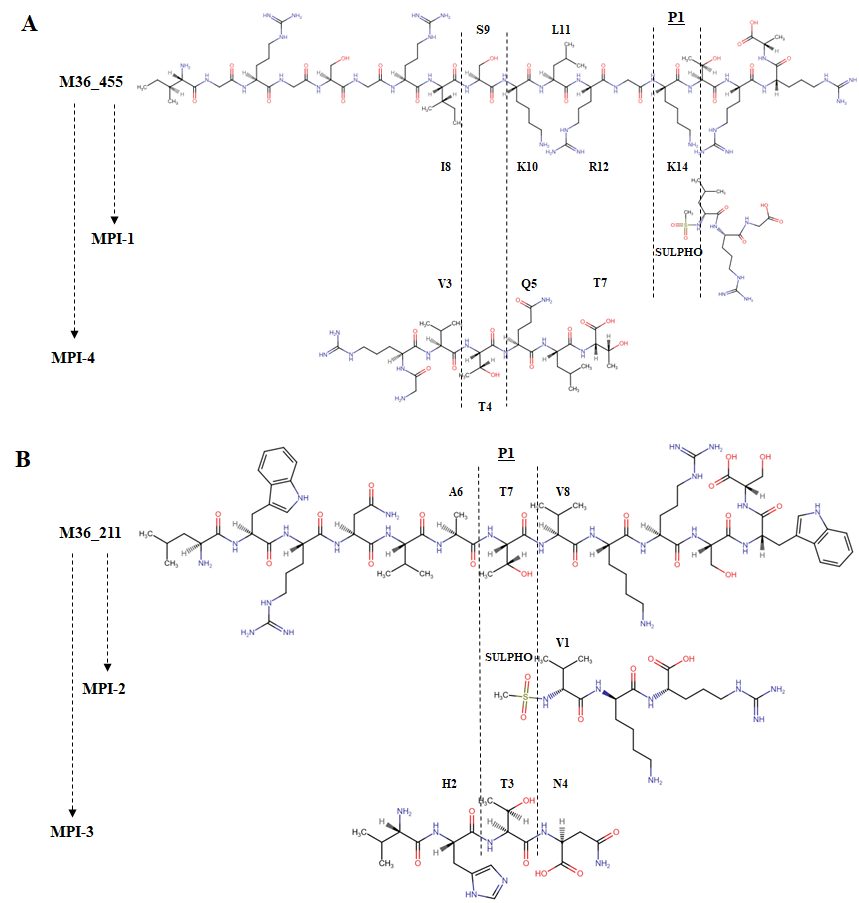

**Figure S5**: Structure based design of inhibitors against CnMpr1. **A**: design of MPI-1 and MPI-4 based on M36_455. **B**: design of MPI-2 and MPI-3 based on M36_211. Labeled amino acids where mutated during the optimization stage. P1 position was highlighted with dashed lines. SULPHO stands for methylsulphonoamide.

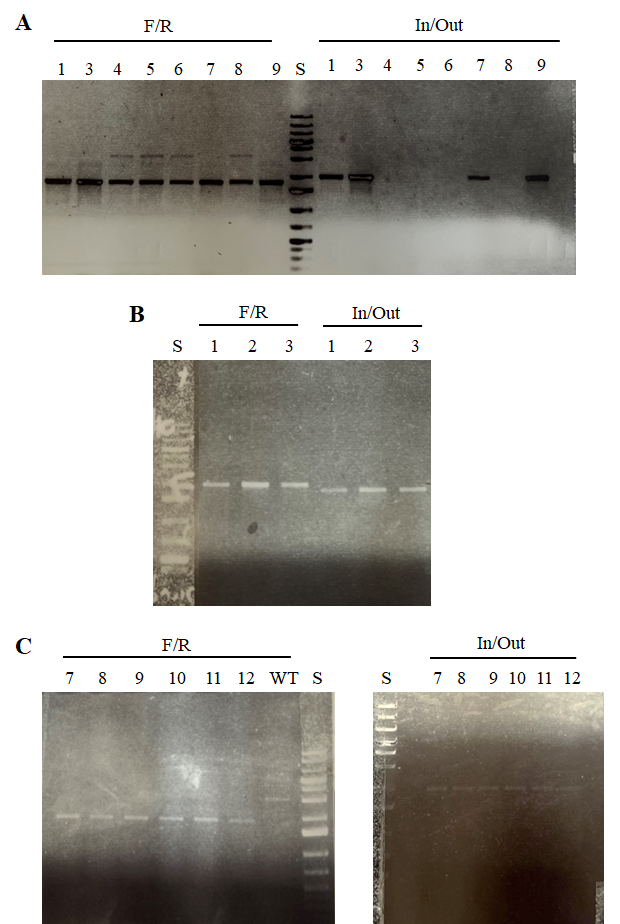

**Figure S6:** Confirmation of gene deletion via PCR on C. neoformans H99 background. **A**: Rim13. **B:** May1. **C:** CnMpr1. F/R stands for forward and reverse PCR. Numbers indicate colonies IDs.

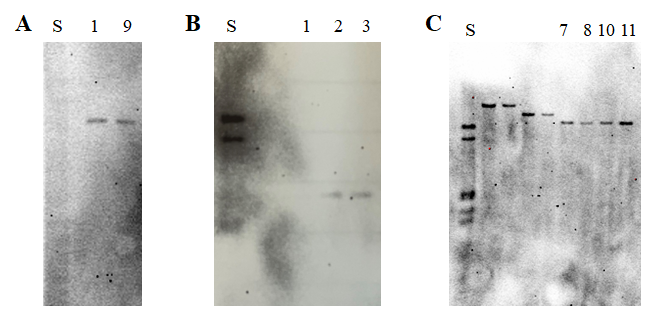

**Figure S7**: **Confirmation of gene deletion via Southern Blot of C. neoformans H99 genome.** **A:** Rim13. **B:** May1. **C:** CnMpr1. F/R stands for forward and reverse PCR. Anti-NAT probe was used for detection.

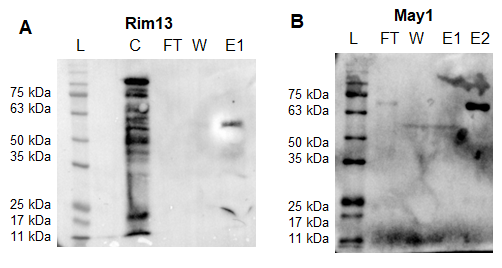

**Figure S8**: **Western blot for peptidase purification**. **A:** Rim13. **B:** May1. L: ladder. C: crude. FT: flow through. W: wash. E: eluates. Anti-FLAG antibody was used for detection.

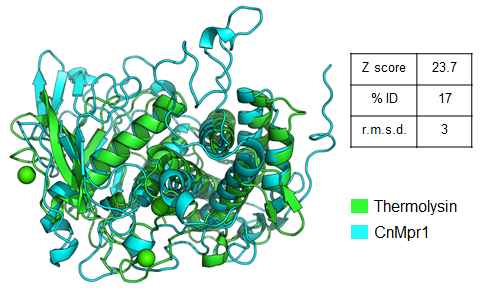

**Figure S9**: **Structural comparison between Thermolysin and CnMpr1**. Z-score indicates structural alignment where higher than 20 indicates homology. % ID reflect sequence alignment. R.M.S.D. stands for root mean square deviation between atoms. Analysis was performed using DALI server (http://ekhidna2.biocenter.helsinki.fi/dali/).

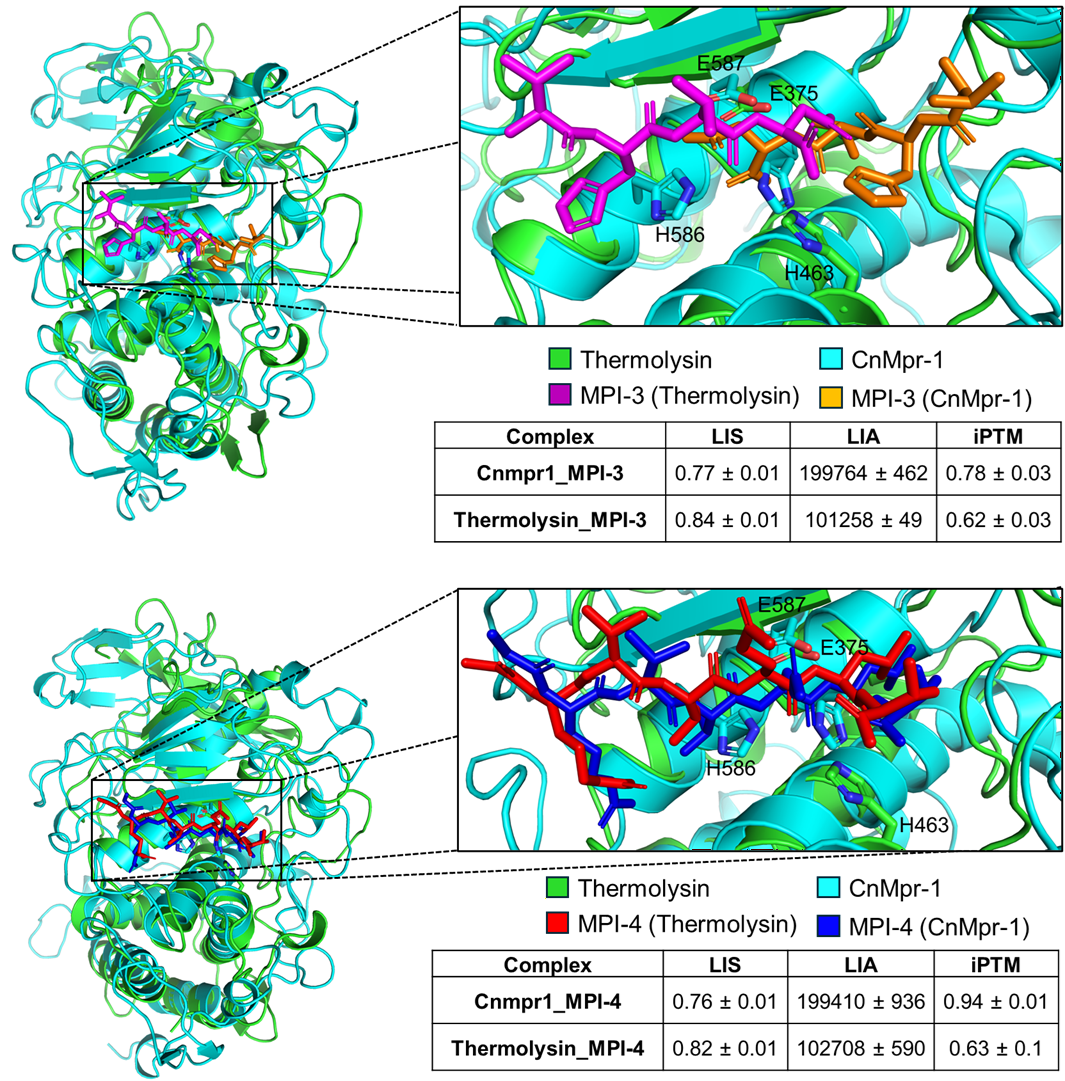

**Figure S10: Prediction of interaction between candidate inhibitors and metalloproteases Thermolysin and CnMpr-1.** **A:** MPI-3. **B:** MPI-4. Thermolysin (Uniprot: P00800) and CnMpr-1 (Uniprot: J9VXZ9) structures in complex with each candidate inhibitor were predicted using AlphaFold-3. Thermolysin carbon backbone is colored green and CnMpr1 cyan. MPI-3 complexed with Thermolysin is colored magenta and with CnMpr-1 orange. MPI-4 complexed with Thermolysin is colored red and with CnMpr-1 dark blue. Structure predictions of proteases were superimposed using the "super" algorithm within PyMol. Interaction scores between the inhibitor candidates and each peptidase were derived from Alpha-Fold-3. Figures were prepared using PyMol.

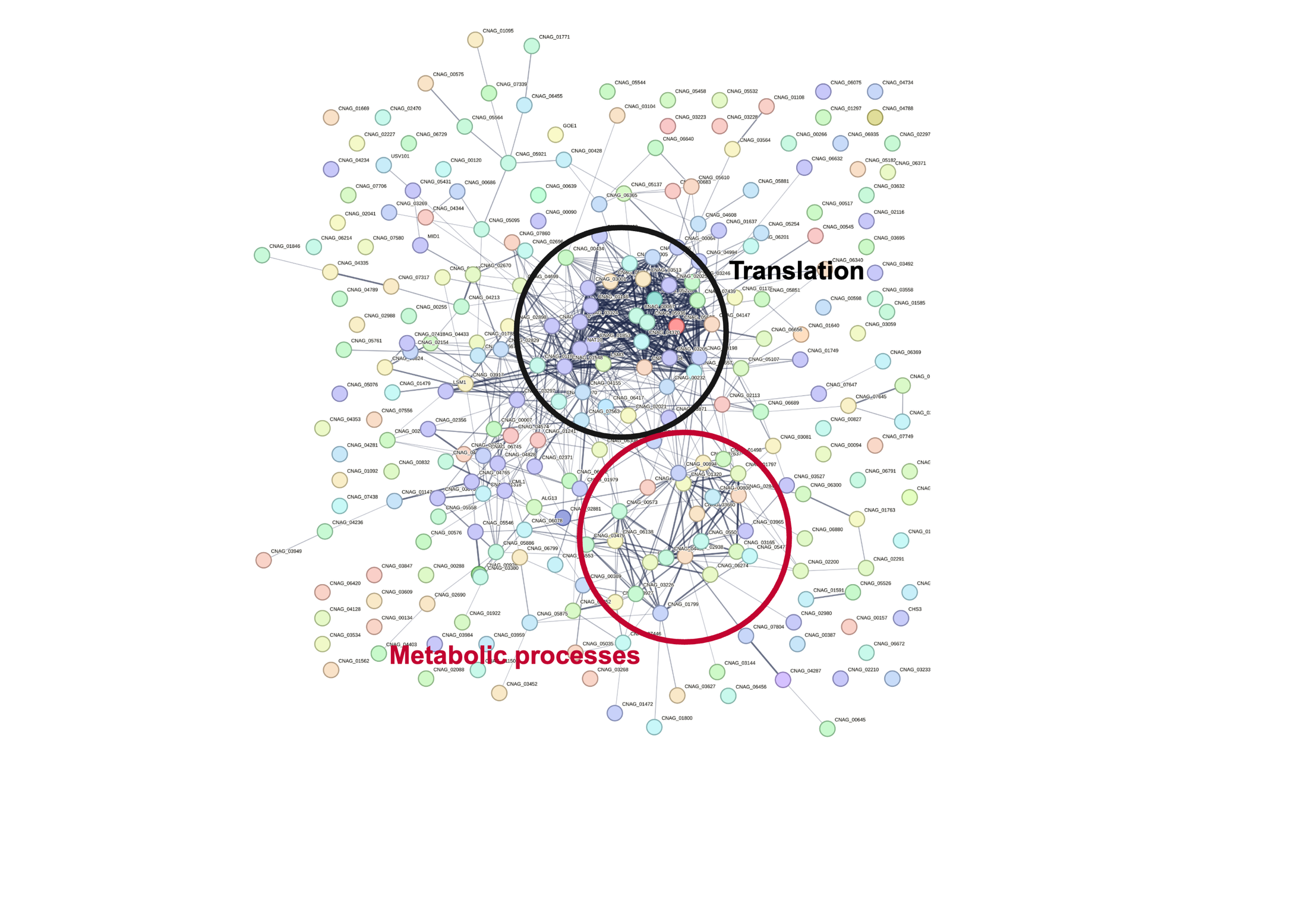

**Figure S11**: **Protein-protein interaction network analysis on proteins with lowered abundance after CPI-1 treatment.** Circles highlight clustered proteins based on known and predicted PPI interactions. Analysis was performed using the STRING web server (https://string-db.org/).
